# Genomic Insights into Multi-Drug-Resistant, *Klebsiella pneumoniae* Isolated from an Infertile Woman: A Comprehensive Analysis of Antibiotic Resistance and Virulence Factors

**DOI:** 10.64898/2026.09.03.749063

**Authors:** Zahid Hasan, Sangita Ahmed, Sabikunnahar Tupur, Mahmuda Yasmin

## Abstract

**Objectives:** *Klebsiella pneumoniae* is a significant global health concern due to its ability to cause diverse infections and its increasing multidrug resistance. This study analyzed the whole-genome sequence of multidrug-resistant *K. pneumoniae* strain CP4, isolated from a vaginal swab of an infertile woman, to characterize its virulence factors, antibiotic resistance mechanisms, and epidemiological features.

**Methods:** The Kirby-Bauer disk diffusion assay was used to screen antibiotic resistance. Whole genome sequencing was conducted using the Illumina MiSeq platform and followed by comprehensive bioinformatic analysis.

**Results:** Antibiotic susceptibility testing revealed that the *K. pneumoniae* strain CP4 exhibited resistance to 11 of 18 tested antibiotics, including carbapenems, and displayed a Multiple Antibiotic Resistance (MAR) index of 0.611. Whole-genome sequencing identified an array of putative virulence and metal-resistant genes, classifying it as a human pathogen with a probability score of 0.887. The strain was characterized as sequence type 327 with K16 capsular and O1/O2v1 somatic antigens. The antibiotic resistome included 38 putative genes conferring resistance to a spectrum of antibiotics. Moreover, point mutations were identified in several genes that ResFinder predicted to be associated with resistance to fluoroquinolones, cephalosporins, and carbapenems. Additionally, *K. pneumoniae* strain CP4 harbored six putative plasmid-replicon types and 454 putative mobile genetic element-associated protein hits among predicted ORFs. Several putative antibiotic resistance genes, including *tet(A)*, *floR*, *mph(A)*, and *aph(6)-Id*, were associated with insertion sequences, transposons, and integrons, indicating a potential for horizontal gene transfer.

**Conclusions:** The genomic features of multidrug-resistant *K. pneumoniae* CP4 highlight its potential for antimicrobial resistance and virulence, emphasizing the need for continued genomic surveillance and targeted interventions.

## 1. Introduction

*K. pneumoniae* is a Gram-negative opportunistic commensal organism that inhabits human mucosal surfaces, including the gastrointestinal tract and oropharynx[1]. It can invade tissues and cause a range of infections, including pneumonia, bloodstream infections, urinary tract infections, and vaginal infections[1,2]. The bacterium is one of the most common causes of both community-acquired and nosocomial infections, often associated with a high fatality rate[1,3].

Aside from its ability to cause various infections, *K. pneumoniae* is of significant concern due to its extensive drug resistance [4]. This bacterium is intrinsically resistant to ampicillin and has acquired resistance to multiple other antibiotics, including carbapenems, beta-lactams, and cephalosporins, attributed mainly to mutation and horizontal gene transfer[3,5]. This has complicated the treatment strategies, resulting in higher mortality rates and a significant economic burden [3]. Considering these features, the World Health Organization has classified *K. pneumoniae* as a critical or high-priority nosocomial pathogen [2].

The challenging drug resistance issue in *K. pneumoniae* is particularly alarming in low-income countries like Bangladesh, where underdeveloped healthcare systems worsen the problem. The emergence of extended-spectrum β-lactamase-producing *K. pneumoniae* (ESBL-KP), including carbapenem-resistant *K. pneumoniae* (CRKP), has been documented in nosocomial infections in Bangladesh, contributing to significant morbidity and mortality [6].

Recent studies have reported the ability of *K. pneumoniae* to interact with human spermatozoa and linked vaginal infections by this pathogen with infertility [7]. By altering sperm motility, increasing the number of necrotic cells, and inducing apoptosis, the *K. pneumoniae* infection may significantly impair fertility [7]. Additionally, *K. pneumoniae* has been implicated in pelvic inflammatory disease (PID) in women of reproductive age [8], a condition that can result in infertility. This issue is especially pressing in Bangladesh, where 15% of women of reproductive age experience infertility, the highest rate among South Asian countries [9]. This further emphasizes the importance of elucidating the underlying mechanism of this opportunistic pathogen’s role in infertility. A recent study by Hasan *et al*., (2025) suggests a possible association between female infertility and opportunistic vaginal pathogens such as *Staphylococcus aureus*, *K. pneumoniae, Enterococcus faecalis,* and *Escherichia coli* [10]. Among these, *K. pneumoniae* remains the least studied concerning female infertility. Also, the prevalence of this organism, especially strains resistant to carbapenems, has become a growing concern in Bangladesh [11].

With this view, to enhance our understanding of the epidemiology and pathogenesis of antibiotic-resistant *K. pneumoniae*, this study characterizes the whole genome sequence of a multidrug-resistant *K. pneumoniae* strain obtained from a vaginal sample of an infertile woman in Bangladesh [10]. Utilizing whole genome sequencing, the study aims to get a comprehensive understanding of the genetic profile, including its antibiotic resistance mechanisms, virulence factors, and develop targeted interventions that alleviate the burden of antibiotic-resistant *K. pneumoniae* infections.

## 2. Material and methods

### 2.1 Isolation and identification of *K. pneumoniae*

Vaginal swabs were obtained by a doctor from infertile women seeking treatment at Bangabandhu Sheikh Mujib Medical University (BSMMU), Bangladesh, following standard protocols [12]. The swabs were inoculated in Brain Heart Infusion (BHI) broth (HiMedia, India), and after 24 hours of enrichment, the samples were streaked onto MacConkey agar (Liofilchem® S.r.l., Italy). Lactose-fermenting pink, gummy colonies were isolated and identified by biochemical tests. Representative *K. pneumoniae* isolates were subjected to *16S rRNA* gene sequencing. Genomic DNA was extracted from 18-hour *K. pneumoniae* culture grown in Luria Bertani (LB) broth (Liofilchem® S.r.l., Italy) at 37 °C using the lab-optimized boiling method[13]. The *16S rRNA* gene was amplified using the universal primers 27F (5′- AGAGTTTGATCCTGGCTCAG-3′) and 1492R (5′-CTACGGCTACCTTGTTACGA-3′) [14], targeting an approximately 1,500 bp region. PCR amplification was performed using Quick-Load® Taq 2X Master Mix (New England Biolabs, Inc., USA) in a Veriti™ 96-Well Thermal Cycler (Applied Biosystems™, USA) with the following lab optimized cycling conditions: initial denaturation at 95 °C for 5 min; 35 cycles of denaturation at 94 °C for 1 min, annealing at 56 °C for 1.5 min, and extension at 72 °C for 1 min; followed by a final extension at 72 °C for 10 min.

The amplified PCR product was subjected to paired-end sequencing using the Illumina platform. Reads were assembled using SeqMan v2.3, and a consensus sequence was generated by aligning overlapping forward and reverse reads. This consensus sequence was then used for taxonomic identification using the Nucleotide BLAST (BLASTn) tool on the NCBI website (https://blast.ncbi.nlm.nih.gov/Blast.cgi). The search was performed against the standard core nucleotide collection (nr/nt) database using the “highly similar sequences (megablast)” option.

### 2.2 Antibiotic sensitivity assay

Antibiotic sensitivity assay was performed following the Kirby-Bauer disk diffusion assay[15]. Approximately 18-hour-old broth cultures grown in LB medium (Liofilchem® S.r.l., Italy) were diluted and adjusted to match the 0.5 McFarland standard. Using a sterile cotton swab, the bacterial suspension was evenly spread over the surface of Mueller-Hinton agar (Liofilchem® S.r.l., Italy) plates. Antibiotic discs (Liofilchem® S.r.l., Italy) were then placed on the inoculated agar, and the plates were incubated at 37°C for 18–24 hours. After incubation, the diameters of the inhibition zones were measured, and the results were interpreted according to the Clinical and Laboratory Standards Institute (CLSI) guidelines[16].

### 2.3 DNA Extraction and Library Preparation for WGS

Based on the antibiotic sensitivity assay, *K. pneumoniae* CP4 was selected for whole genome analysis. For whole genome analysis, genomic DNA was extracted using the Qiagen DNeasy Blood & Tissue Kit (250) (Qiagen, #69504). Following evaluation of DNA concentration using Nanodrop (Thermo Fisher Scientific, ND-1000), the DNA was sent to the International Centre for Diarrheal Disease Research, Bangladesh (icddrb) for WGS. For library preparation, the Illumina DNA Prep Reagent Kit and an automated liquid handler (epMotion 5075) were utilized. The prepared DNA libraries were sequenced using the Illumina MiSeq platform with paired-end sequencing. Sequencing was performed using a 2 × 250 bp read configuration.

### 2.4 Whole Genome Sequence Analysis

#### 2.4.1 Quality Assessment, Trimming, and Genome Assembly

The quality of the raw data was assessed using FastQC v0.12.1 (https://www.bioinformatics.babraham.ac.uk/projects/fastqc/). Subsequently, fastp v0.25.0 [17] was employed for adapter removal and quality trimming with the following parameters: trimming of 20 bases from the 5′ ends of both forward and reverse reads (-f 20 -F 20), trimming of 83 bases from the 3′ ends (-t 83 -T 83), and discarding reads shorter than 80 bp after trimming (-l 80). The fastp command was as follows: fastp -i input_R1.fastq.gz -I input_R2.fastq.gz -o output_R1.fastq.gz -O output_R2.fastq.gz -f 20 -F 20 -t 83 -T 83 -l 80 -j fastp.json. These parameters were chosen based on FastQC results. The fastp summary showed mean read lengths of 240 and 241 bp for the forward and reverse reads, respectively, before quality filtering and trimming. Following trimming and quality filtering, the mean read length was 145 bp. After confirming the quality of the raw reads, de novo assembly of the trimmed paired-end sequences was performed using SPAdes v3.13.1 [18]. Contigs shorter than 200 bp were excluded from the final assembly. The resulting draft genome assembly was then assessed for its quality employing QUAST v5.3.0 [19], CheckM v1.2.3[20] and completeness using BUSCO 6.0.0 [21].

#### 2.4.2 Genome Annotation, Pathways, and Metabolite Analysis

Annotation of the draft genome was carried out using prokka 1.14.6 [22] and visualized using Proksee version 1.1.3[23], while subsystem-based rapid annotation was conducted using the RAST [24]. The NCBI Prokaryotic Genome Annotation Pipeline (PGAP) version 6.7 was also used to annotate the draft genome (https://www.ncbi.nlm.nih.gov/genome/annotation_prok/). Additionally, a circular genome map was generated using PanExplorer[25] to visualize gene arrangements. Functional characterization of the genome was conducted using the BlastKOALA annotation server[26] and eggNOG-mapper v2[27]. Using the KEGG Orthology (KO) numbers assigned to the genes by BlastKOALA, the KEGG Mapper Reconstruct tool was employed to map the genes against the KEGG Pathway database for functional interpretation of the genome.

The assembled genome was analyzed for secondary metabolite biosynthesis gene clusters using the antiSMASH bacterial version[28].

#### 2.4.3 Pathway Enrichment Analysis

Following functional annotation with eggNOG-mapper v2, pathway enrichment analysis was performed using EC (Enzyme Commission) numbers and gene-preferred names in ShinyGo 0.80[29]. *Klebsiella pneumoniae* (taxonomy ID: 573) was selected as the reference organism. As no custom background gene set was specified, all protein-coding genes in the selected *K. pneumoniae* genome were used as the default background for the enrichment analysis. A total of 836 functional categories were assessed, and categories with an FDR-adjusted p-value < 0.05 were considered significantly overrepresented. The analysis was performed to identify functional categories that were statistically overrepresented in the CP4 genome relative to the reference gene set.

#### 2.4.4 Detection of Virulence and Metal Resistance Genes

The presence of virulence factors and metal resistance genes in the isolate were identified using the BacWGSTdb 2.0 [30] and BacMet 2.0 [31], respectively.

#### 2.4.5 Detection of Resistance Genes, Mutations, and Mobile Genetic Elements

ABRicate v1.0.1 (https://github.com/tseemann/abricate) was employed to identify antibiotic-resistant genes present in the draft genome. Analyses were performed against three databases using the default thresholds: CARD database version 2025-Jan-14 [32] (Threshold for minimum %identity: 80%; %coverage: 90%) to capture a broader set of potential resistance genes, ResFinder [33] (Threshold for minimum %identity: 95%; %coverage: 95%), and NCBI database version 2025-Jan-14 [34] (Threshold for minimum %identity: 95%; %coverage: 95%) to focus on high-confidence hits. ResFinder 4.5.0 [35] was used to identify chromosomal point mutations in relevant antimicrobial resistance genes.

Putative plasmid replicon sequence types in the draft genome assembly were identified using ABRicate v1.0.1 with the PlasmidFinder database (version 2025-Jan-14) [36] (Threshold for minimum %identity: 80%; %coverage: 90%). The matched replicon gene regions were extracted from the corresponding contigs and visualized using SnapGene® software v7.2.1 (from Dotmatics; available at https://www.snapgene.com/). Other mobile genetic elements (MGE) associated protein hits were determined using mobileOG-db (Beatrix 1.6) [37] and VRprofile2 [38] was employed to determine the antibiotic-resistance genes carried by the MGEs. Probable candidate Horizontal Gene Transfer (HGT) events were identified using Alien Hunter 1.7 [39]. All the genome maps were generated using Proksee version 1.1.3 [23].

#### 2.4.6 Detection of Prophage region and CRISPR-Cas Systems

Phigaro 2.3.0[40] and PHASTEST 3.0[41] were used to identify prophage regions, while CRISPRCasFinder 4.2.20[42] was employed to detect CRISPR-Cas systems in the *K. pneumoniae* CP4 genome.

#### 2.4.7 Sequence typing, Serotyping, and Pathogenicity Prediction

Multilocus Sequence Typing (MLST) and determination of O- and K-serotypes for *K. pneumoniae*, based on lipopolysaccharide (LPS, O-antigen) and capsular polysaccharide (K-antigen), were conducted using the Pathogenwatch Platform v22.3.8 (https://pathogen.watch/). Additionally, Pathogenfinder [43] was utilized to determine if the isolate was a probable human pathogen.

#### 2.4.8 Phylogenetic and Pangenome Analysis

The Similar Genome Finder tool of the BV-BRC web service (https://www.bv-brc.org/) was used to identify genomes similar to *K. pneumoniae* CP4, the sequenced isolate. Using the accession numbers obtained from the tool, 48 similar whole genome sequences from diverse sources and regions were downloaded from NCBI GenBank. These sequences were utilized to perform SNP-based phylogenetic analysis using CSI Phylogeny [44], which applies stringent quality control filters to ensure statistical robustness. The genome assemblies were provided in FASTA format, with *K. pneumoniae* CP4 used as the reference genome. The analysis employed the following parameters: a minimum depth of 10 at SNP positions, a relative depth threshold of 10, a minimum distance of 10 bp between SNPs (to prune closely spaced variants), a minimum SNP quality score of 30, a minimum read mapping quality of 25, and a Z-score threshold of 1.96, corresponding to a 95% confidence interval. Positions that did not meet the specified quality and depth criteria were excluded from SNP analysis. The resulting SNP alignment comprised 95,054 variable positions across the analyzed genomes. The proportion of valid positions relative to the CP4 reference ranged from 88.40% to 97.47% among the individual genomes (Supplementary Table 10). An unrooted phylogenetic tree was generated from the high-confidence SNP alignment, with branch-support values (0–1) retained in the resulting tree. The tree was subsequently visualized using iTOL v6 in a circular layout [45]. The Roary pangenome pipeline, available in IPGA v1.09[46] (Integrated Prokaryotic Genome and Pan-Genome Analysis service) and PanExplorer[25], was employed for the pangenome of the study isolate *K. pneumoniae* CP4, along with 48 sequences downloaded from NCBI GenBank. Roary analysis was performed using a minimum BLASTp identity threshold of 70%, with a core-gene ratio of 0.95 and a support value of −1. Specifically, IPGA v1.09 was used to generate the Average Nucleotide Identity (ANI) heatmap and core/pangenome rarefaction curves, while PanExplorer was used to visualize the distribution of core, strain-specific, and dispensable genes, COG categories, and Circos plots illustrating genomic arrangement. The cluster-sharing dendrogram showing relationships among genomes was generated using IPGA v1.09.

### 2.5 Data Availability

The Whole Genome Shotgun project has been deposited at DDBJ/ENA/GenBank (https://www.ncbi.nlm.nih.gov/genbank) under the accession JBDPJK000000000, Bioproject: PRJNA1107790, Biosample: SAMN41212664, SRA: SRR29031371. The *16S rRNA* gene sequence has also been deposited to GenBank (https://www.ncbi.nlm.nih.gov/genbank) under Accession No. PP345570. The genome annotation data generated using Prokka & RAST and genome quality assessment data by Quast are available on Zenodo [47].

### 2.6 Ethics and Consent

Ethical approval for this study was obtained from the Faculty of Biological Sciences, University of Dhaka (Approval No. 258/Biol.Scs; approved on 25 September 2024). Sample collection was conducted with the permission of the ethical committee, chaired by Professor Dr. Md. Akhter Hossain Khan. Written informed consent was obtained from all participants prior to the collection of swab samples. Participants were informed about the objectives of the study and the intended publication of the findings. The study was conducted in accordance with applicable local laws and ethical guidelines.

## 3. Results

A total of 16 *K. pneumoniae* were isolated and identified from 55 vaginal swabs of infertile women (Supplementary Table 1). Initial screening for antibiotic resistance of these isolates against eight antibiotics from different classes revealed that one isolate, *K. pneumoniae* CP4, was resistant to all the antibiotics except ciprofloxacin (Intermediate resistance) (Supplementary Table 1). Based on this resistance profile, *K. pneumoniae* CP4 was selected for further investigation and subsequently tested against an additional 10 antibiotics. It was found to be resistant to 4 of these, resulting in resistance to 11 out of the 18 antibiotics tested against the selected *K. pneumoniae* CP4 isolate (Table 1). Resistance was observed to representatives of different classes of antibiotics (Table 1), including carbapenems (meropenem and doripenem), monobactam (aztreonam), penicillins (ampicillin, amoxicillin-clavulanate), third generation cephalosporin (ceftazidime), macrolides (azithromycin and erythromycin), and the 2nd generation aminoglycoside (gentamycin), fosfomycin and chloramphenicol. The isolate showed intermediate resistance to fluoroquinolone (ciprofloxacin), 1st generation tetracycline (tetracycline), and was sensitive to 2nd generation tetracycline (doxycycline), and the 3rd generation aminoglycoside (netilmicin) (Table 1).

**Table 1:**
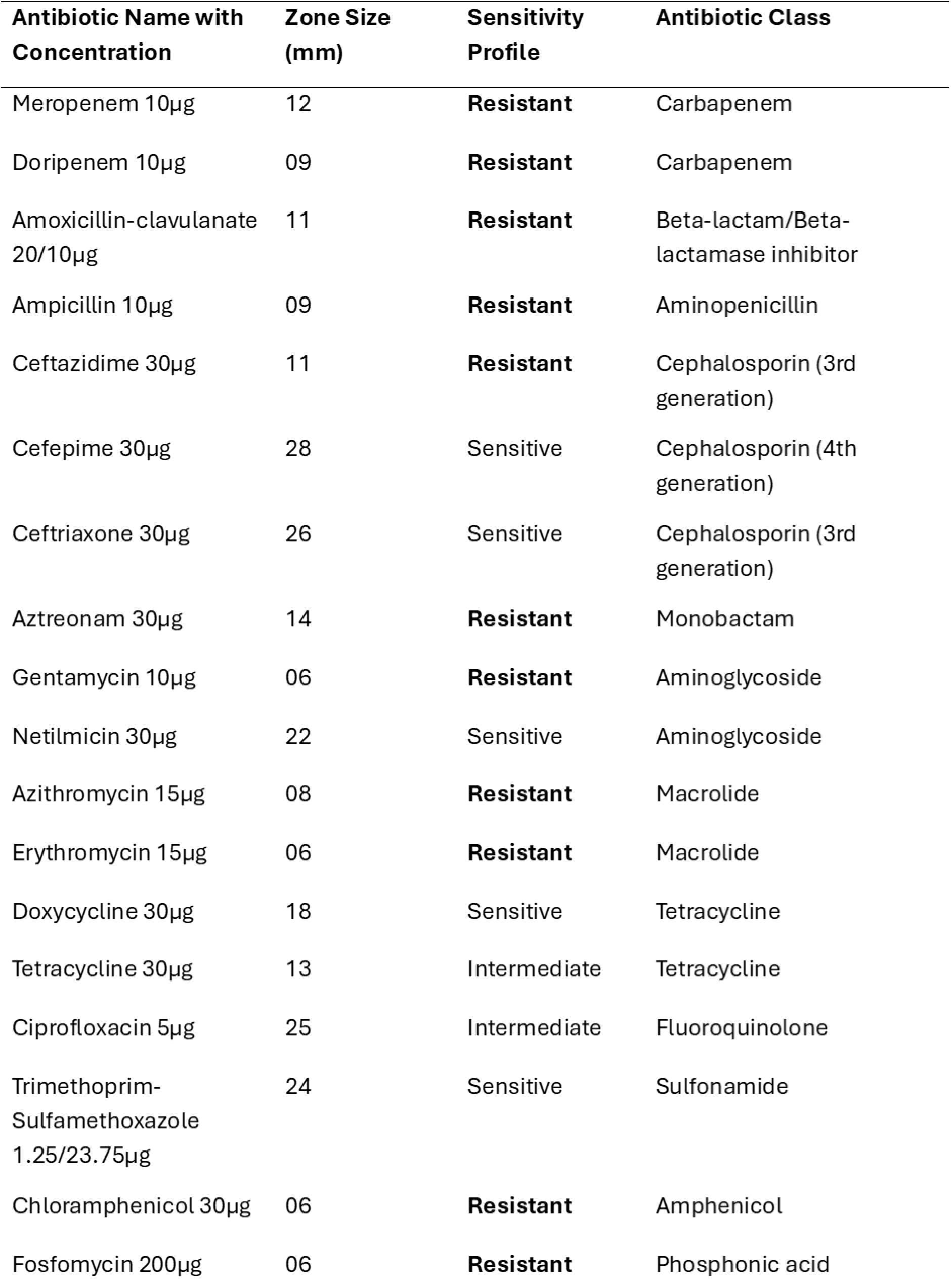
Antibiotic Resistance Profile of *K. pneumoniae* CP4 by Kirby-Bauer Disk Diffusion Assay.

This *K. pneumoniae* CP4 was obtained from a 32-year-old female patient diagnosed with primary infertility and pelvic inflammatory disease (PID), who sought infertility treatment at Bangladesh Medical University, Dhaka. This isolate, obtained from a woman with infertility-related complications, displaying resistance to 11 out of 18 different antibiotics and exhibiting a MAR index of 0.611, was selected for whole-genome sequencing to gain insight into the genomic basis of multidrug resistance and potential genomic basis contributing to vaginal infection and infertility.

### 3.1 Quality Assessment of WGS

An Illumina sequencing run produced a total of 875,323 paired-end reads. Following trimming, 801,723 paired-end reads were utilized in the assembly process, and the resulting draft genome achieved 98.6% completeness with no mismatches. The L50 and N50 values were 4 and 547,596, respectively, with a GC content of 57.23% (Supplementary Table 2). The assembled genome shared 99.19% Average Nucleotide Identity (ANI) with *K. pneumoniae* subsp. *pneumoniae* (Supplementary Table 2).

### 3.2 Genome Annotation and Functional Analysis

Genome annotation was performed using Prokka 1.14.6 (Figure 1A), complemented by subsystem annotation via the RAST (Figure 1B), revealing a genome length of 5,378,058 bp (Supplementary Table 3). Prokka identified 4,997 coding sequences (CDS) and 5,090 genes, along with 10 rRNA and 82 tRNA (Supplementary Table 3). RAST analysis identified a total of 390 subsystems (Supplementary Table 3), spanning various functional categories such as stress response, dormancy, sporulation, virulence, metal resistance, and antibiotic resistance (Figure 1B). The annotation data are available on Zenodo[47]. The Circos genome map (Supplementary Figure 1) of *K. pneumoniae* CP4 displays the arrangement of genes transcribed in both forward and reverse directions, along with core genes and strain-specific genes.

**Figure 1:**
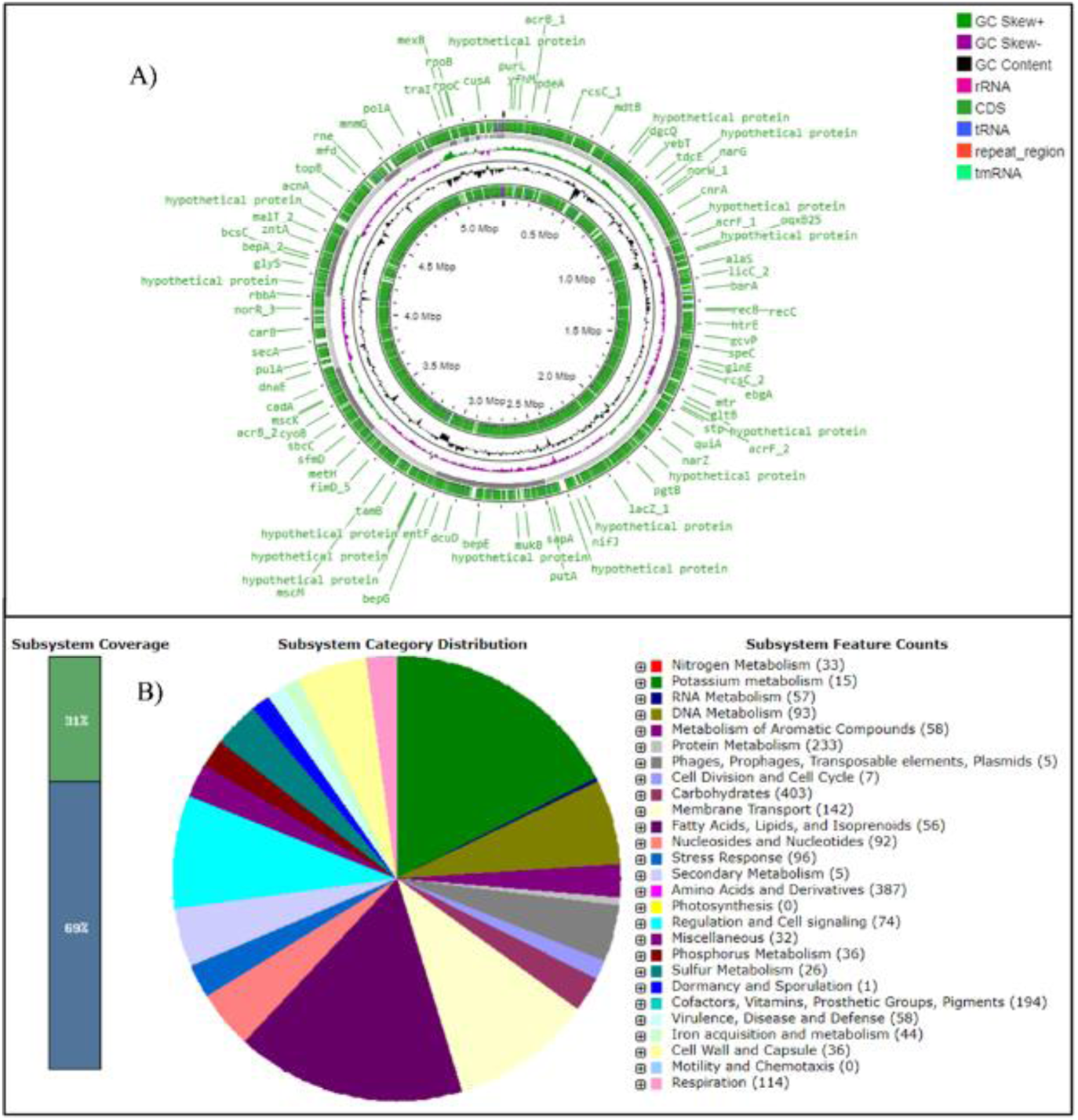
**A)** Genome map generated by Proksee based on annotation performed with Prokka, displaying coding sequences (CDS), rRNA, tRNA, GC content, GC skew, repeat regions, and other genomic features. **B)** Rapid genome annotation with RAST using subsystem technology, including feature counts for various functional categories.

Functional annotation of the genome by BlastKOALA revealed that the major functional pathways include signaling and cellular processes, genetic information processing, environmental information processing (Figure 2A).

**Figure 2:**
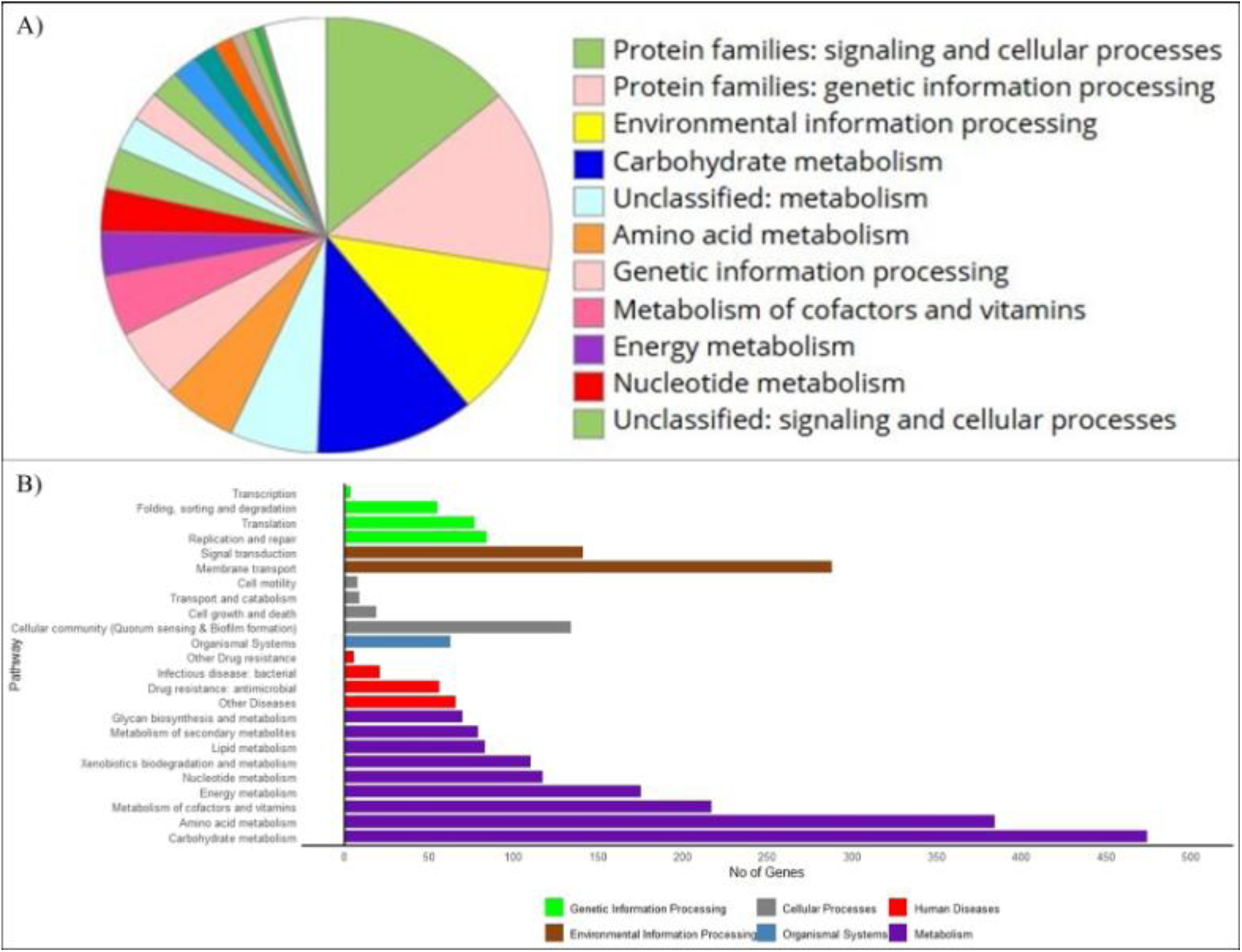
**A)** Functional annotation of the genome using BlastKOALA, highlighting the distribution of functional categories. **B)** KEGG pathway mapping of *K. pneumoniae* CP4 using the KEGG Mapper Reconstruct Tool

KEGG pathway mapping analysis of *K. pneumoniae* CP4 revealed that the majority of its biochemical pathways were associated with carbohydrate metabolism, amino acid metabolism, cofactor and vitamin metabolism, energy metabolism, membrane transport, and signal transduction (Figure 2B).

Furthermore, four secondary metabolite biosynthesis gene clusters (Supplementary Figure 2) were identified by antiSMASH in the study isolate, including redox-cofactor, thiopeptide, NRP-metallophore, and RiPP-like clusters.

### 3.3 Pathway Enrichment

The pathway enrichment analysis identified biological processes that were significantly overrepresented in the *K. pneumoniae* CP4 genome relative to the reference gene set. Oxidoreductase activity displayed the highest overrepresentation, followed by catabolism of organic substances and other metabolic activities, indicating significant representation of catalytic and metabolic functions (Supplementary Figure 3A). Ribonucleoprotein and ribosomal processes also showed significant overrepresentation, representing functional categories associated with genetic information processing (Supplementary Figure 3B). Additionally, biosynthetic processes, including cellular biosynthetic processes and organonitrogen compound biosynthesis, were significantly overrepresented, indicating the representation of metabolic and biosynthetic functions among the annotated CP4 genes.

### 3.4 Virulence Profile

Whole genome sequencing revealed the presence of an array of putative virulence factors (69 putative virulence factors) in *K. pneumoniae* CP4, particularly those associated with adhesion to human mucosal or epithelial surfaces, biofilm formation, tissue invasion, immunomodulation, and iron acquisition (Supplementary Table 4). Notably, genes encoding Type VI secretion systems (T6SS), which are crucial for bactericidal activity, and genes for type 1 fimbriae involved in epithelial cell adhesion and invasion, were identified. Genes for type 3 fimbriae, which mediate biofilm formation on both biotic and abiotic surfaces, were also present. The presence of enterotoxin genes was also observed in the isolate (Supplementary Table 4). Additionally, genes encoding enterobactin, a siderophore critical for iron acquisition, overcoming host defenses, and facilitating penetration into deeper tissues, were identified. The detection of all these virulence genes emphasizes the pathogenic potential of *K. pneumoniae* CP4.

### 3.5 Antibiotic resistance profile

Screening the genome sequence using the ABRicate tool version 1.0.1 against the CARD database identified a total of 38 putative antimicrobial resistance (AMR) genes, while 20 putative AMR genes were found using the NCBI (with 100% coverage for all genes) and ResFinder databases (Figure 3A). These genes confer resistance against antibiotics such as Carbapenem, Beta-lactams, Chloramphenicol, Cephalosporin, Macrolide, Tetracycline, Sulfonamide, Streptomycin, Trimethoprim, Quinolone, Rifamycin, Amikacin, Kanamycin, Tobramycin, Gentamicin, Fosfomycin, etc. (Supplementary Table 5).

**Figure 3:**
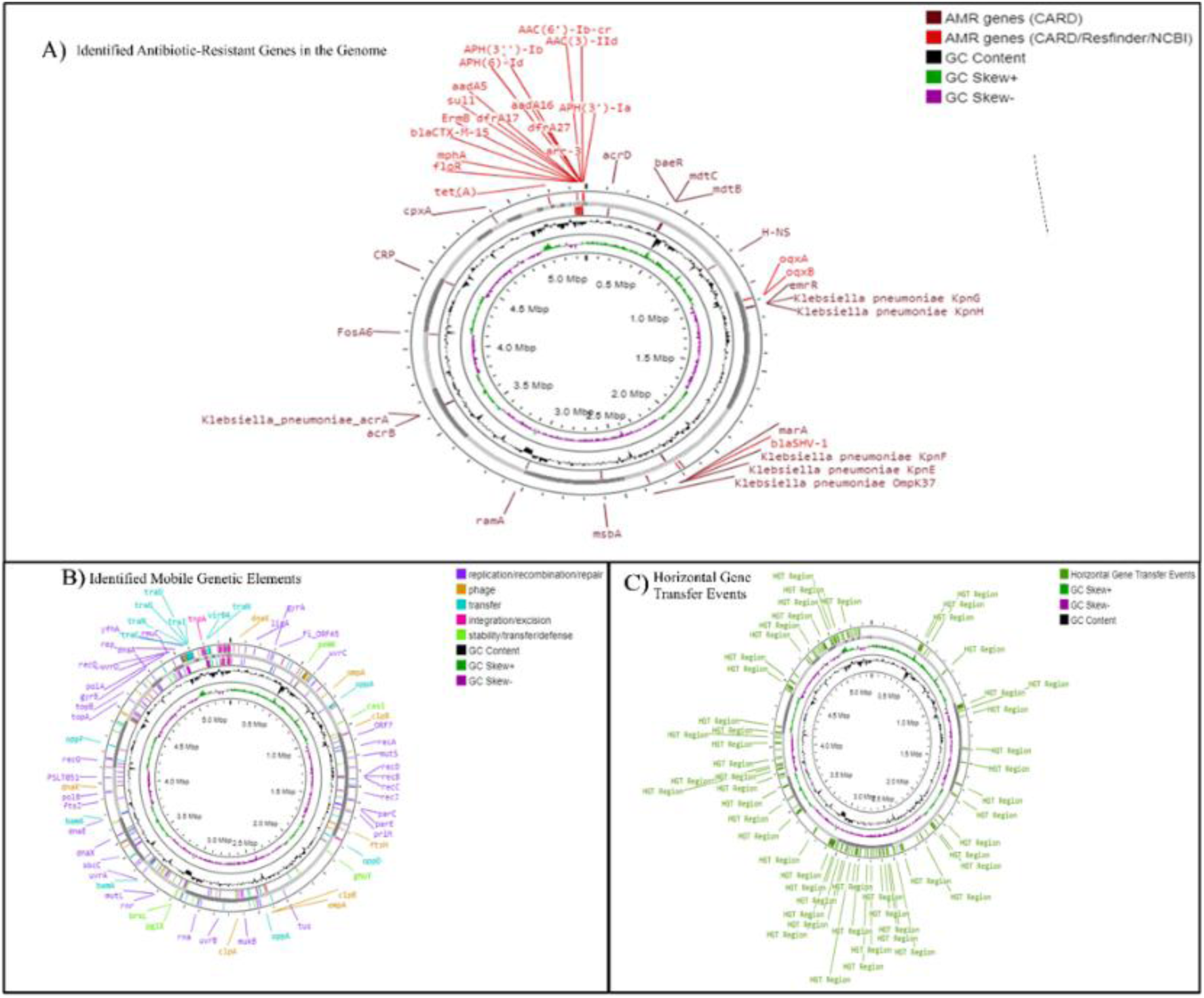
Genome Map Highlighting **A)** The Antibiotic-Resistant Genes Identified by ABRicate 1.0.1 Using CARD, ResFinder, and NCBI Databases. Genes detected by the NCBI, ResFinder, and CARD databases are depicted in red, while those detected only by the CARD database are shown in maroon **B)** The Presence of Mobile Genetic Elements in the *K. pneumoniae* CP4 Genome, identified by MobileOG-db. **C)** The Probable Horizontal Gene Transfer Events Identified by Alien Hunter. The genome maps were Generated Using Proksee.

Particularly, the detection of two beta-lactamases, *bla*_SHV-101_ and *bla*_CTX-M-15_, one of the most prevalent enzymes in ESBL isolates, validates the resistance of this isolate to beta-lactam antibiotics. The genome analysis also revealed the presence of several putative efflux pumps associated with antibiotic resistance, including *tet(A)*, a tetracycline efflux pump; *acrD*, an aminoglycoside efflux pump; and *oqxA* and *oqxB*, components of the RND efflux pump OqxAB, conferring resistance to fluoroquinolones. The *acrA* and *acrB* components of the acrAB-TolC efflux pump and *ramA*, a positive regulator that increases the expression of the AcrAB operon, were also identified. Additionally, the *floR* gene, a chloramphenicol exporter, as well as the *mphA* and *fosA6* genes, which confer resistance to macrolides and fosfomycin, respectively, were detected (Figure 3A)[32].

*K. pneumoniae KpnE* and *KpnF*, components of the small multidrug resistance (SMR) family antibiotic efflux pump KpnEF, and *KpnG* and *KpnH*, components of the major facilitator superfamily (MFS) antibiotic efflux pump KpnG-KpnH, were identified in the genome. These efflux pumps confer resistance to a broad spectrum of antibiotics, including cephalosporins, carbapenems, aminoglycosides, and macrolides (Figure 3A)[32].

### 3.6 Mutations Associated with Antibiotic Resistance

ResFinder 4.5.0 detected several chromosomal point mutations in the genes *AcrR*, which encodes a repressor of the acrAB efflux pump operon system, and in the genes *OmpK36* and *OmpK37,* which encode outer membrane porins. According to the ResFinder predictions, these mutations were associated with resistance to fluoroquinolones, cephalosporins, and carbapenems (Table 2).

**Table 2:** Identified chromosomal point mutations in Antimicrobial resistant genes of *K. pneumoniae* CP4 using ResFinder 4.5.0 and their associated antimicrobial resistance.

| **Mutated Gene** | **Nucleotide Substitution** | **Amino Acid Substitutions and Position** | **Conferred Resistance** |
| --- | --- | --- | --- |
| *acrR* | CCG -> CGG | P161R | Fluoroquinolone |
| *acrR* | GGC -> GCC | G164A | Fluoroquinolone |
| *acrR* | TTC -> TCC | F172S | Fluoroquinolone |
| *acrR* | CGA -> GGG | R173G | Fluoroquinolone |
| *acrR* | CTC -> GTC | L195V | Fluoroquinolone |
| *acrR* | TTT -> ATT | F197I | Fluoroquinolone |
| *acrR* | AAG -> ATG | K201M | Fluoroquinolone |
| *ompK36* | AAC -> AGC | N49S | Cephalosporins |
| *ompK36* | CTT -> GTA | L59V | Cephalosporins |
| *ompK36* | CTG -> AGC | L191S | Cephalosporins |
| *ompK36* | TTC -> TGG | F207W | Cephalosporins |
| *ompK36* | GCT -> TCT | A217S | Carbapenem |
| *ompK36* | AAC -> CAC | N218H | Carbapenem |
| *ompK36* | GAT -> GAG | D224E | Cephalosporins |
| *ompK36* | CTG -> GTT | L228V | Cephalosporins |
| *ompK36* | GAA -> CGT | E232R | Cephalosporins |
| *ompK36* | ACT -> TCT | T254S | Cephalosporins |
| *ompK37* | ATT -> ATG | I70M | Carbapenem |
| *ompK37* | ATT -> ATG | I128M | Carbapenem |

### 3.7 Metal and Chemical Resistance

Putative Metal and chemical-resistant genes were also identified in the isolate, particularly those conferring resistance to copper, manganese, nickel, cobalt, arsenic, tungsten, molybdenum. Genes associated with resistance to n-hexane, p-xylene, and acid tolerance were also evident in the genome (Supplementary Table 6).

### 3.8 Mobile Genetic Elements and Horizontal Gene Transfer

The PlasmidFinder database identified six putative plasmid-replicon type sequences in the *K. pneumoniae* CP4 draft genome assembly (Supplementary Table 7). Two putative plasmid-replicon types belonged to the IncF plasmid family (IncFIB(AP001918)_1 and IncFII(29)_1_pUTI89), three to Col-type plasmids (Col156_1, ColRNAI_1 and Col440II_1), and one to the IncR incompatibility group (IncR_1). Putative plasmid-replicon type sequences showing >95% identity to reference plasmids were extracted and visualized using SnapGene Viewer v7.2.1 (Supplementary Figure 4). No virulence or antimicrobial resistance genes were detected within the identified putative plasmid-replicon type sequences. A hypothetical protein was detected within the IncFIB(AP001918)_1-associated contig region (Supplementary Figure 4A). Several restriction enzyme recognition sites were also observed within the analyzed sequences (Supplementary Figure 4).

MobileOG-db identified 454 putative mobile genetic element-associated protein hits among predicted ORFs in the genome, comprising 90 integration/excision, 145 replication/recombination/repair, 62 phage-related, 53 stability/transfer/defense, and 104 transfer-associated hits (Figure 3B). Several antibiotic resistance genes such as *tet(A), floR, mph(A), aph(6)-Id* were carried within insertion sequences, transposons, and integrons (Supplementary Table 8). Additionally, Alien Hunter identified 96 candidate horizontally acquired regions in the genome (Figure 3C).

### 3.9 Prophage Regions and CRISPR-Cas Systems

Two prophage regions comprising 89 putative prophage genes were identified in *the K. pneumoniae* CP4 genome using Phigaro (Figure 4A). In contrast, PHASTEST analysis detected two putative intact prophage regions (37,241bp & 33,191bp) containing a total of 105 phage-related genes. Additionally, five CRISPR arrays, eight Cas genes, and one Cas gene cluster were identified in the genome (Figure 4B).

**Figure 4:**
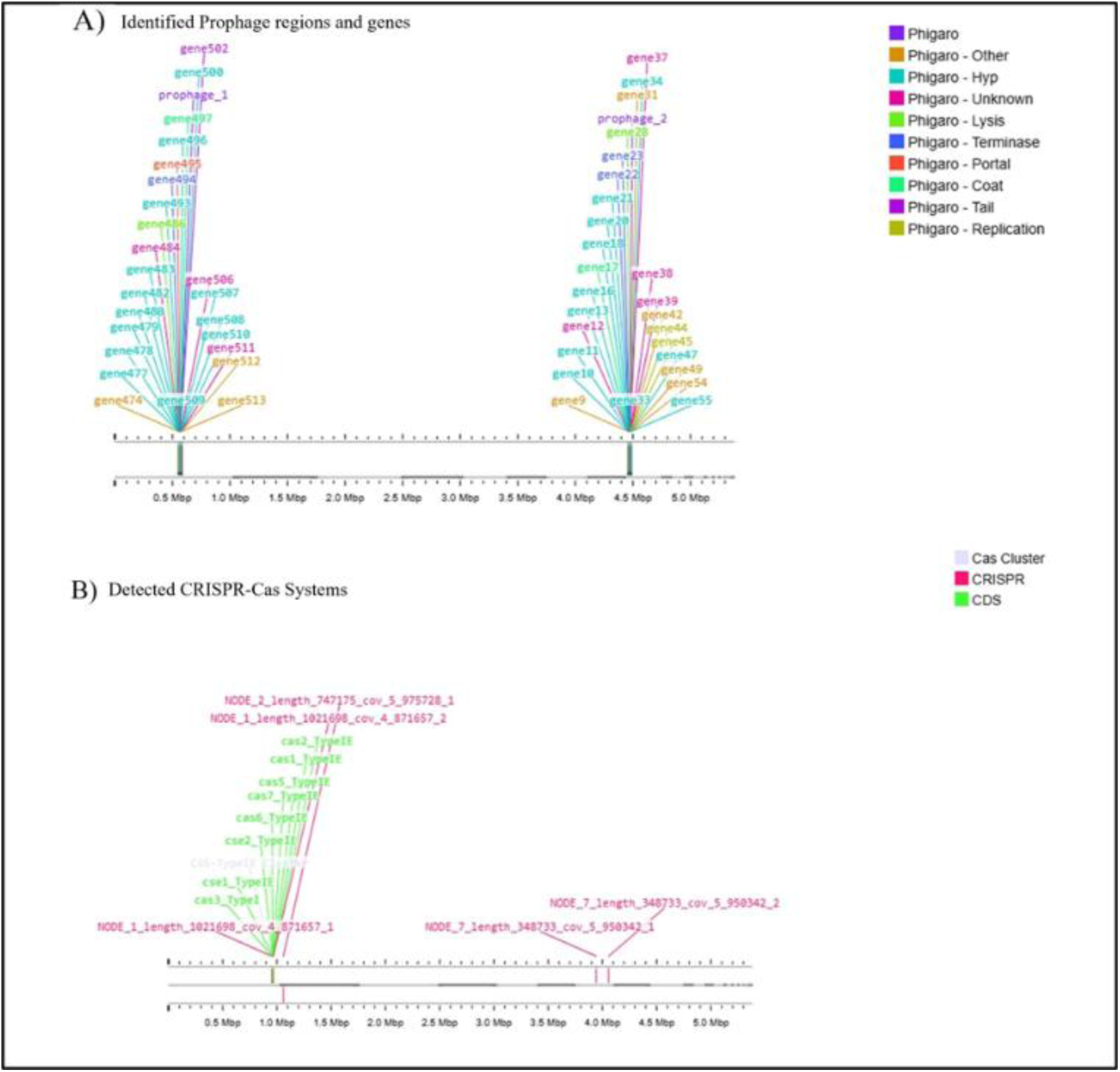
Illustration showing **A)** Prophage regions and genes detected using Phigaro 2.3.0, and **B)** CRISPR-Cas systems detected using CRISPRCasFinder4.2.20 in the *K. pneumoniae* CP4 genome.

### 3.10 Isolate Typing and Pathogenicity

The allelic profile for the seven housekeeping genes *gapA, infB, mdh, pgi, phoE, rpoB*, and *tonB* retrieved from the Pathogenwatch server was 2-1-1-1-10-1-19, classifying *K. pneumoniae* CP4 as sequence type 327 (ST327). The isolate was identified as the K16 serotype based on its capsular polysaccharide (K-antigen), while lipopolysaccharide (LPS) O-antigen classified it as O1/O2v1 serotype. Pathogenfinder predicted this isolate as a human pathogen with a probability score of 0.887, matching 340 pathogenic families and 18 non-pathogenic families.

### 3.11 Phylogenetic and Pangenome Analysis

SNP-based phylogenetic analysis revealed that isolates of the same sequence type (ST) exhibit the highest sequence similarity and cluster together on the same branch of the phylogenetic tree. The query isolate, *K. pneumoniae* CP4, which belongs to ST327, is closely related to the ST327 isolate GCA_025909935.1 (Figure 5A), isolated from Cantonese cabbage in China (Supplementary Table 9). These two isolates shared over 97.4% valid genome positions, supporting their high genetic similarity (Supplementary Table 10). These two ST327 isolates are more closely related to the ST13 isolate GCA_018275295.1, derived from a human stool sample in China, than to other ST327 isolates (Figure 5A). This suggests potential sequence similarity between ST327 and ST13. The phylogenetic clustering indicates that isolates of the same ST tend to group together, but inter-ST sequence similarities may occur, possibly influenced by the isolation source and geographic region. The Average Nucleotide Identity (ANI) analysis (Figure 5B) and cluster share analysis (Supplementary Figure 5) demonstrated that *K. pneumoniae* CP4 exhibited the closest relatedness to isolate GCA_025909935.1, which was obtained from Cantonese cabbage in China, consistent with the findings from the phylogenetic analysis. *K. pneumoniae* strain CP4 and GCA_025909935.1, exhibit an Average Nucleotide Identity (ANI) of approximately 99%, as inferred from the dark brown shading on the ANI heatmap and the associated color scale (95–100%) (Figure 5B).

**Figure 5:**
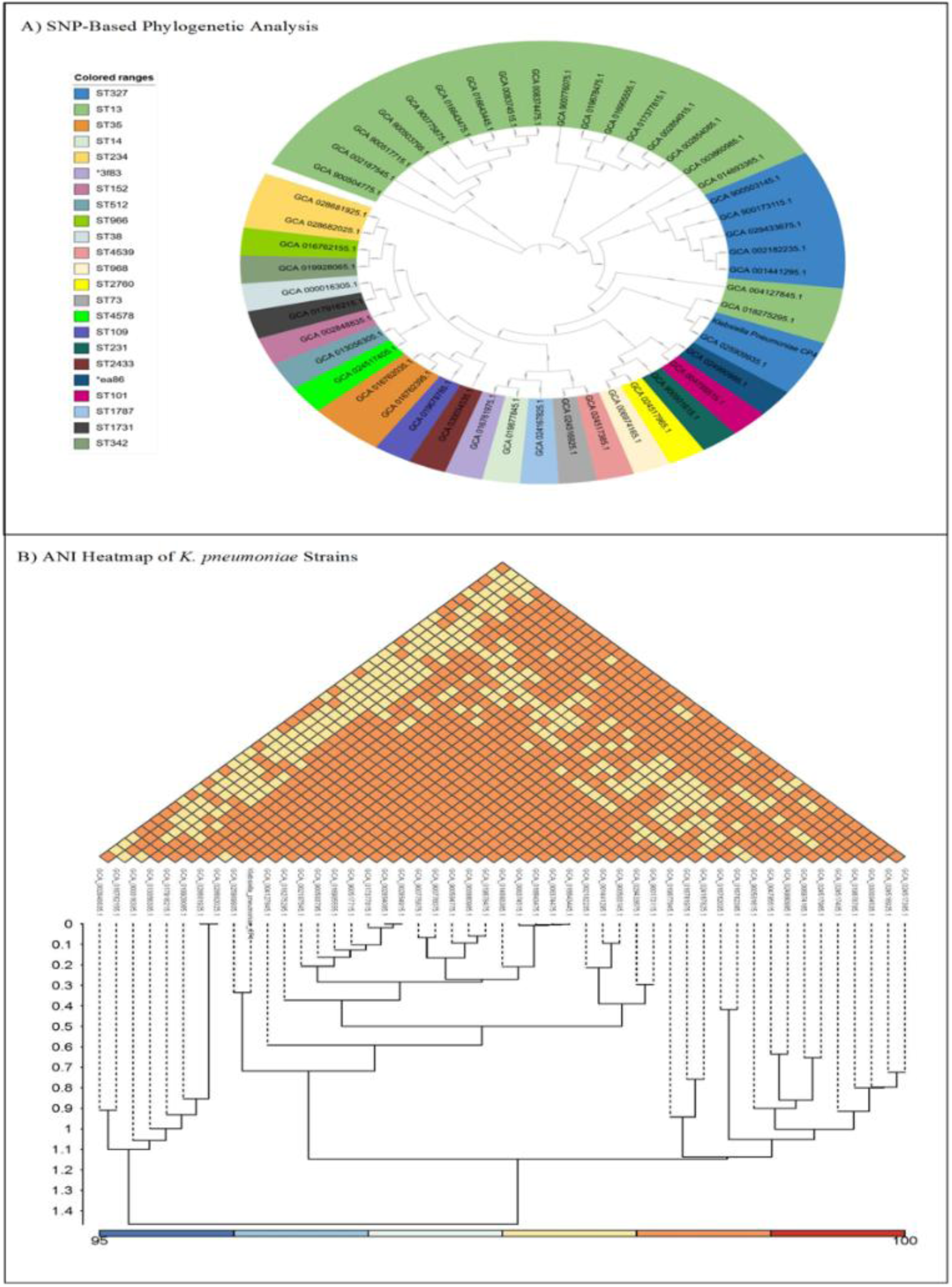
**A)** SNP-Based Phylogenetic Analysis of *K. pneumoniae* CP4. The phylogenetic tree was constructed using CSI Phylogeny with genomes of various *K. pneumoniae* isolates downloaded from NCBI GenBank and visualized using iTOL, **B)** Pangenome analysis of *K. pneumoniae* using IPGA v1.09: Average Nucleotide Identity (ANI) heatmap

Roary based pan-genome analysis of 48 *K. pneumoniae* genomes downloaded from NCBI GenBank, along with the study isolate *K. pneumoniae* CP4 (Supplementary Table 9) revealed 12,718 clusters, consisting of 2,452 (19.3%) core genes, 3,694 (29%) strain-specific genes, and 6,572 (51.7%) dispensable genes (Figure 6A). The distribution of COG (Clusters of Orthologous Groups) categories across the *K. pneumoniae* genomes indicated that, similar to other strains, most of the genes in *K. pneumoniae* CP4 are involved in information storage and processing (Figure 6C). The core-pan rarefaction curve shows that as the number of genomes increases; the number of gene clusters increases. However, the number of core gene clusters initially decreases drastically and then flattens, suggesting a saturation point where the addition of new genomes contributes minimally to the discovery of new core gene clusters (Figure 6B).

**Figure 6:**
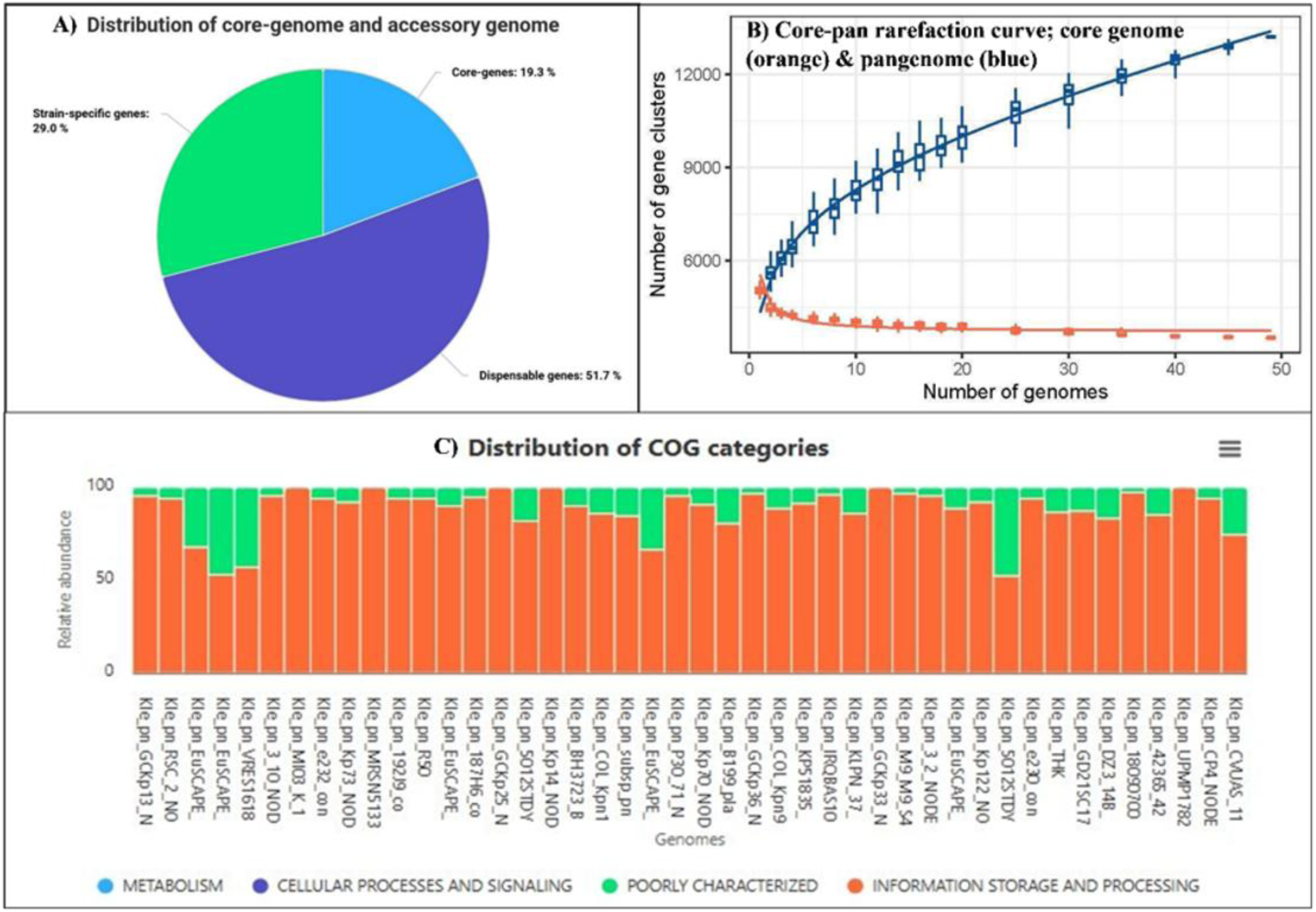
Pangenome analysis of *K. pneumoniae* CP4 using **A)** PanExplorer, showing the distribution of core genes, strain-speciic genes, and dispensable genes, **B)** IPGA v1.09, core genome (orange) and pangenome (blue) rarefaction curves, **C)** PanExplorer, illustrating the distribution of COG (clusters of Orthologous Groups) categories.

## 3. Discussion

This study focuses on characterizing the whole genome sequence of a multidrug-resistant (MDR) *K. pneumoniae* isolate obtained from the vaginal swab of an infertile patient with pelvic inflammatory disease (PID) in Bangladesh. Comprehensive analysis of the genomic features aims to provide insights into virulence factors, resistance determinants, guide treatment strategies, and support effective management of MDR infections.

Pathway enrichment analysis of *K. pneumoniae* CP4 revealed significant overrepresentation of carbohydrate metabolism, other metabolic pathways, and various biosynthetic processes. Bacterial metabolism plays a crucial role in mediating cellular responses to antibiotic treatment [48]. Several studies have demonstrated that carbohydrate metabolism contributes significantly to biofilm formation and maintenance[48]. However, the overrepresentation of these metabolic categories in CP4 does not by itself establish their direct contribution to virulence, antibiotic resistance, or biofilm-associated survival. Additionally, oxidoreductase activity was among the most highly overrepresented functional categories identified by the enrichment analysis. Oxidoreductases participate in diverse cellular redox reactions and can contribute to cellular responses to oxidative stress [49]. The oxidoreductase activity is also involved in iron homeostasis, that supports growth and virulence in the host[49]. Nevertheless, the overrepresentation of oxidoreductase-related genes in CP4 does not by itself demonstrate enhanced oxidative-stress resistance, detoxification of host-derived reactive oxygen species. Further comparative or expression-based studies would be required to determine whether these functions contribute to specific phenotypes in CP4.

Genome analysis revealed that *K. pneumoniae* CP4 contains major virulence factor genes commonly reported in this pathogen, including those encoding capsule, siderophores, lipopolysaccharide (LPS), fimbriae, outer membrane proteins, Type VI secretion system, and enterotoxin [50]. Notably, the strain harbors putative genes associated with the K16 capsular type and an O1-associated LPS O-antigen locus (O1/O2v1), which have been reported among some hypervirulent *K. pneumoniae* strains. These features may indicate the presence of virulence-associated characteristics in *K. pneumoniae* CP4; however, additional phenotypic and genomic evidence is required to classify the strain as hypervirulent [51–53]. In addition, multiple metal resistance genes were identified, which may enhance survival in harsh, toxic environments and facilitate the acquisition of antimicrobial resistance genes[54].

The identified putative secondary metabolite biosynthetic gene clusters also contribute to metal acquisition, virulence, antimicrobial activity, and other bioactive properties in the bacterium. Notably, the thiopeptide identified in this bacterium may provide a competitive advantage over other bacteria due to its potent antibacterial activity and ability to overcome antibiotic resistance, particularly against Gram-positive bacteria[55]. Together these features underscore the virulent nature of *K. pneumoniae* CP4, and support the strain’s classification as a human pathogen, as predicted by Pathogenfinder.

The challenges extend beyond virulence factors, as genome analysis of the *K. pneumoniae* CP4 revealed 38 putative antimicrobial resistance (AMR) genes along with several mutations in AMR genes, conferring resistance to major classes of antibiotics. The phenotypic expression of these AMR genes was also confirmed by disk diffusion assay. A major concern is the isolate’s resistance to carbapenems, which are regarded as drugs of last resort[56]. However, no carbapenemase genes were detected in the genome. The observed carbapenem resistance may therefore have putative genetic determinants, including alterations in outer membrane porins such as OmpK36 and OmpK37, which have been associated with reduced susceptibility to β-lactams, including carbapenems [32]. Multiple chromosomal point mutations were identified in *ompK36* and *ompK37*. However, their specific contribution to the observed resistance phenotype could not be established in the present study. Nevertheless, previous studies have reported that alterations or loss of major outer membrane porins, particularly OmpK36, can contribute to increased resistance to carbapenems and cephalosporins [32,57–59]. Recent reports have also described an association between production of CTX-M-15 enzyme and carbapenem resistance in non-carbapenemase-producing *K. pneumoniae*, particularly when combined with alterations in outer membrane porins [58,60]. Therefore, from a genomic perspective, the combination of alterations in *ompK36* and *ompK37* and the presence of *bla_CTX-M-15_* may represent a possible genomic explanation for the observed carbapenem resistance in *K. pneumoniae* CP4. Recently, the *bla*_CTX-M-15_ gene has been detected in the highest prevalence in clinical *K. pneumoniae* isolates collected from Dhaka[61].

Furthermore, the *K. pneumoniae* CP4 genome displays its dynamic ability to acquire and disseminate antibiotic resistance genes through mobile genetic elements and horizontal gene transfer [5]. Six putative plasmid-replicon types were detected, including IncFIB, IncFII, ColRNAI_1, and Col440II_1. These are associated with carbapenem resistance in XDR *K. pneumoniae*. Moreover, IncF plasmids are linked to virulence in *Enterobacteriaceae*, including *K. pneumoniae* [62]. In particular, the IncFIB plasmid has been reported to disseminate the *bla*_NDM_, *bla*_SHV_, *bla*_CTX−M_, and *bla*_OXA_ genes in *K. pneumoniae* [63]. However, the present study did not reconstruct or circularize the plasmid sequences. Therefore, the association of these resistance or virulence determinants with the identified replicons in CP4 could not be established.

Putative antibiotic resistance genes were also identified within insertion sequences, transposons, and integrons, possibly obtained from other bacteria via horizontal transfer. Identification of multiple candidate horizontally acquired regions by Alien Hunter suggests the potential occurrence of horizontal gene transfer in the genome.

Type I-E CRISPR-Cas system, the most common CRISPR‒Cas system reported in the *K. pneumoniae,* was identified in the *K. pneumoniae* CP4 genome. This system is known to limit the acquisition of external genetic elements, including plasmids and bacteriophages, and is generally associated with reduced antibiotic resistance due to its restriction of Antibiotic-Resistant Gene (ARG)-bearing mobile genetic elements [64]. However, despite its defensive role, CRISPR-Cas systems often co-exist with numerous ARGs, as seen in the CP4 genome. This apparent contradiction may be due to the strong selective pressure for ARG acquisition, and phages expressing anti-CRISPR proteins [64]. Recently, Li *et al*., demonstrated a link between the type I- E* CRISPR-Cas system in *K. pneumoniae* with virulence [65]. The isolates containing this CRISPR-Cas system were moderate or strong biofilm producers and had a higher frequency of virulence genes [66]. Any link between the presence of CRISPR-Cas system in the *K. pneumoniae* CP4 with virulence and antibiotic resistance needs further investigation.

Two intact prophages belonging to the Siphoviridae family were also identified in the CP4 genome. Prophages are known to contribute to the genome plasticity of *K. pneumoniae*, facilitating acquisition and chromosomal integration of antibiotic resistance genes [67]. Phages can mediate ARG transfer through generalized, specialized, or auto-transduction, contributing to the spread of antimicrobial resistance [68]. Prophages can also enhance bacterial survival under stress conditions, including antibiotic exposure, by modulating host metabolism and stress responses [68]. Therefore, the prophages detected in the *K. pneumoniae* CP4 genome might have some contribution to its virulence and antibiotic resistance.

The phylogenetic analysis positioned the study isolate within the same clade as an ST327 *K. pneumoniae* isolate recovered from Cantonese cabbage in China and an ST13 isolate obtained from a stool sample in China. However, as per the literature, there is no report of ST327 *K. pneumoniae* in Bangladesh so far, suggesting a possible emergence of new sequence types of *K. pneumoniae* in Bangladesh. Pangenome analysis of the study isolate alongside 48 other *K. pneumoniae* isolates revealed genetic diversity among the species from different regions.

In our previous study, we observed a potential association between *K. pneumoniae* and female infertility based on the analysis of vaginal samples from infertile women [10]. The isolates examined in the present study, including *K. pneumoniae* CP4, were obtained from these vaginal samples. Given the limited understanding of the potential role of *K. pneumoniae* in the reproductive tract, we investigated genomic features of CP4 that may provide hypotheses for future studies. However, a definite mechanism cannot be concluded from genomic data, which needs further comprehensive experimental data.

The genome of *K. pneumoniae* CP4 contains genes encoding type 1 and type 3 fimbriae, which are established virulence-associated structures involved in adhesion and colonization in *K. pneumoniae* [69]. Their presence in CP4 indicates the genetic potential for these functions; however, their contribution to vaginal colonization or interaction with reproductive-tract cells was not investigated in the present study.

Additionally, studies have reported effects of *E. coli* lipopolysaccharide (LPS) on sperm viability and DNA integrity [70]. Although *K. pneumoniae* CP4 possesses genes required for LPS biosynthesis, the present study provides no experimental evidence that CP4 LPS affects sperm viability or DNA integrity.

Recent studies have found *Klebsiella* in abundance in the cervix and endometrial samples of endometriosis patients compared to the control group, suggesting the possibility that inflammation triggered by this bacterium might have been attributed to endometriosis, another major factor in female infertility [71].

The genome of *K. pneumoniae* CP4 contains genes encoding several virulence-associated factors, including LPS, enterobactin, and capsular polysaccharide. These features indicate the genetic potential for functions associated with bacterial survival and virulence; however, their contribution to infertility cannot be established from the present genomic data. Further studies using multiple clinical isolates and appropriate in vitro and in vivo models are required to determine whether *K. pneumoniae* has a role in reproductive health.

## 4. Conclusion

Multidrug-resistant isolates, particularly those classified as critical pathogens by the WHO, such as *K. pneumoniae*, pose a significant threat to global public health. In this study, we employed comprehensive whole-genome sequencing to characterize a multidrug-resistant vaginal isolate of *K. pneumoniae* obtained from an infertile woman.

This study presents an extensive genomic characterization of *K. pneumoniae* CP4, covering functional and metabolic profiling, virulence and metal resistance genes, mobile genetic elements involved in gene dissemination, and traits that facilitate infection and pathogen survival. Phylogenetic analysis also revealed its relatedness to other global isolates. The genome harbors numerous putative virulence factors, antimicrobial resistance genes, and other features that support its classification as a pathogen of clinical concern.

While the findings underscore the importance of genome-based surveillance and the urgent need for improved antibiotic stewardship and novel therapeutics, it is important to note that the conclusions, such as the isolate’s pathogenic potential, are based solely on in silico predictions. A major limitation of this study is the absence of experimental validation and phenotypic characterization, which will be the focus of future research to confirm the functional implications of the genomic findings.

## Declaration of competing interest

The authors declare that they have no known competing financial interests or personal relationships that could have appeared to influence the work reported in this paper.

## Supplementary Materials

**Supplementary Table 1:** Identification of 16 *Klebsiella pneumoniae* isolates from vaginal swabs of 55 infertile women using biochemical tests and *16S rDNA* sequencing (GenBank Accession Numbers provided). Primary screening of antibiotic-resistant isolates was conducted using eight antibiotics to select candidates for whole-genome sequencing.

| Isolates | AUG | ATM | CAZ | C | CN | CIP | FOS | MRP | Identification / Gene Bank Accession |
| --- | --- | --- | --- | --- | --- | --- | --- | --- | --- |
| 1 | S | R | R | I | I | S | R | I | Biochemical tests |
| 2 | I | I | R | S | S | I | R | R | Biochemical tests |
| 3 | I | S | S | S | S | I | I | R | Biochemical tests |
| **CP4** | **R** | **R** | **R** | **R** | **R** | **I** | **R** | **R** | **PP345570** |
| 5 | I | I | R | S | I | S | R | R | Biochemical test |
| 6 | R | S | R | R | R | R | R | S | PP345572 |
| 7 | R | S | S | S | I | I | S | R | PP345573 |
| 8 | I | I | R | S | I | I | I | R | PP345575 |
| 9 | I | S | R | S | S | S | R | I | Biochemical tests |
| 10 | I | S | I | S | S | S | S | R | Biochemical tests |
| 11 | I | I | R | S | S | I | R | R | Biochemical tests |
| 12 | I | I | R | S | I | S | R | R | PP346490 |
| 13 | I | S | R | S | S | S | R | R | Biochemical tests |
| 14 | I | I | R | S | I | I | R | R | PP346491 |
| 15 | I | S | R | S | I | I | R | R | Biochemical tests |
| 16 | I | R | R | S | S | I | R | I | Biochemical tests |

\*\*AUG- Amoxicillin + clavulanate, ATM-Aztreonam, CAZ- Ceftazidime, C-Chloramphenicol, FOS- Fosfomycin, CN- Gentamycin, CIP- Ciprofloxacin, MRP- meropenem, R-Resistant, I-Intermediate, S-Sensitive

**Supplementary Table 2:** General Features of the Assembled Genome of *K. pneumoniae* CP4.

| Features | Data for *K. pneumoniae* CP4 | Tools used |
| --- | --- | --- |
| Number of Paired-end Reads after trimming | 801,723 | FastQC v0.12.1 |
| Assembly Length (bp) | 5378058 | Quast 5.3.0 |
| Number of contigs | 61 | Quast 5.3.0 |
| GC content (%) | 57.27 | Quast 5.3.0 |
| Largest contig (bp) | 1,021,698 | Quast 5.3.0 |
| N50 | 547,596 | Quast 5.3.0 |
| N90 | 101,234 | Quast 5.3.0 |
| L50 | 4 | Quast 5.3.0 |
| L90 | 10 | Quast 5.3.0 |
| N's per 100Kbp | 0.00 | Quast 5.3.0 |
| N'S | 0.00 | Quast 5.3.0 |
| Genome completeness (%) | 98.6 | BUSCO 6.0.0 |
|  | 98.42 | CheckM v1.2.3 |
| Genome coverage | 7.16x | PGAP |
| ANI (%) | 99.19 | CheckM v1.2.3 |

**Supplementary Table 3:**
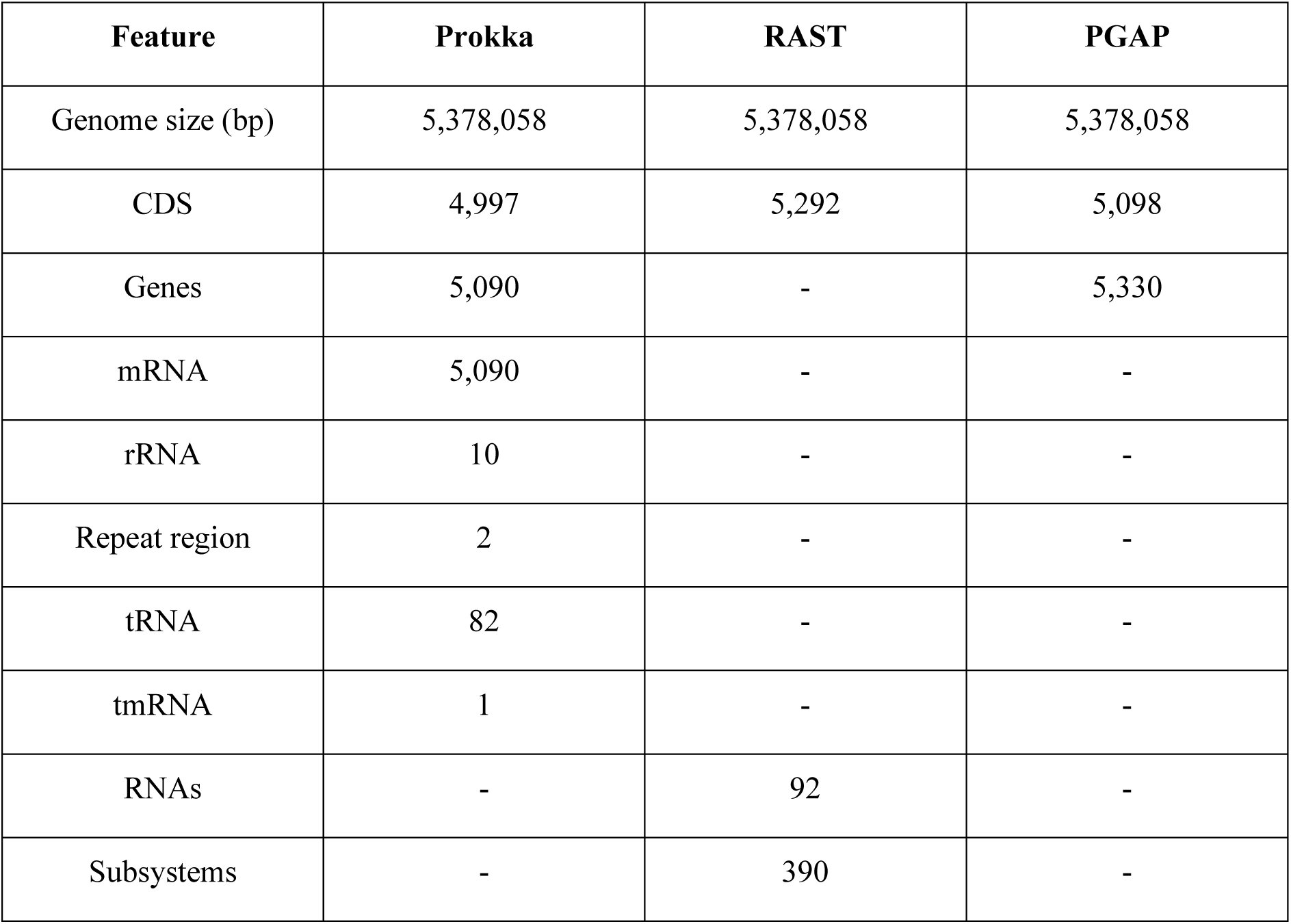
General Features of *K. pneumoniae* CP4 Genome Annotation Using Prokka, RAST and Prokaryotic Genome Annotation Pipeline (PGAP)

**Supplementary Table 4:**
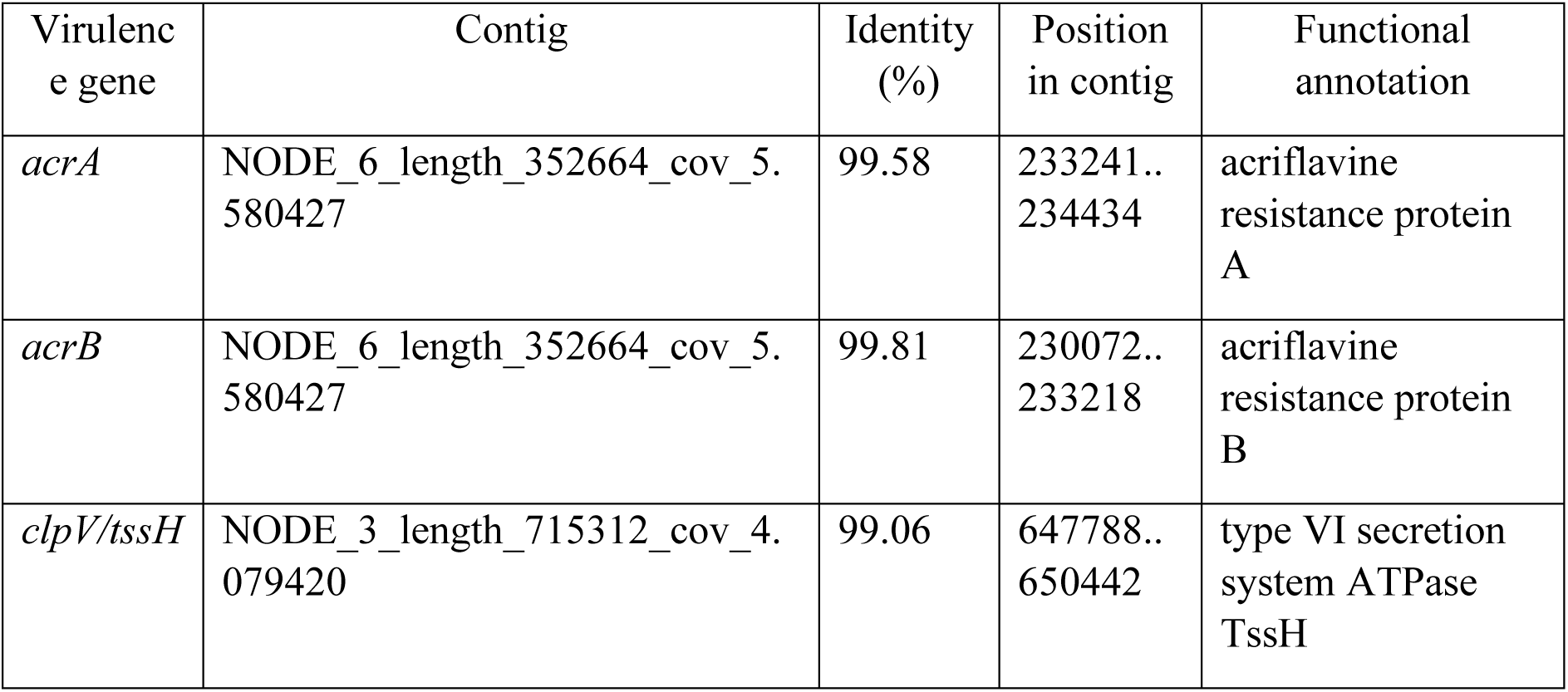
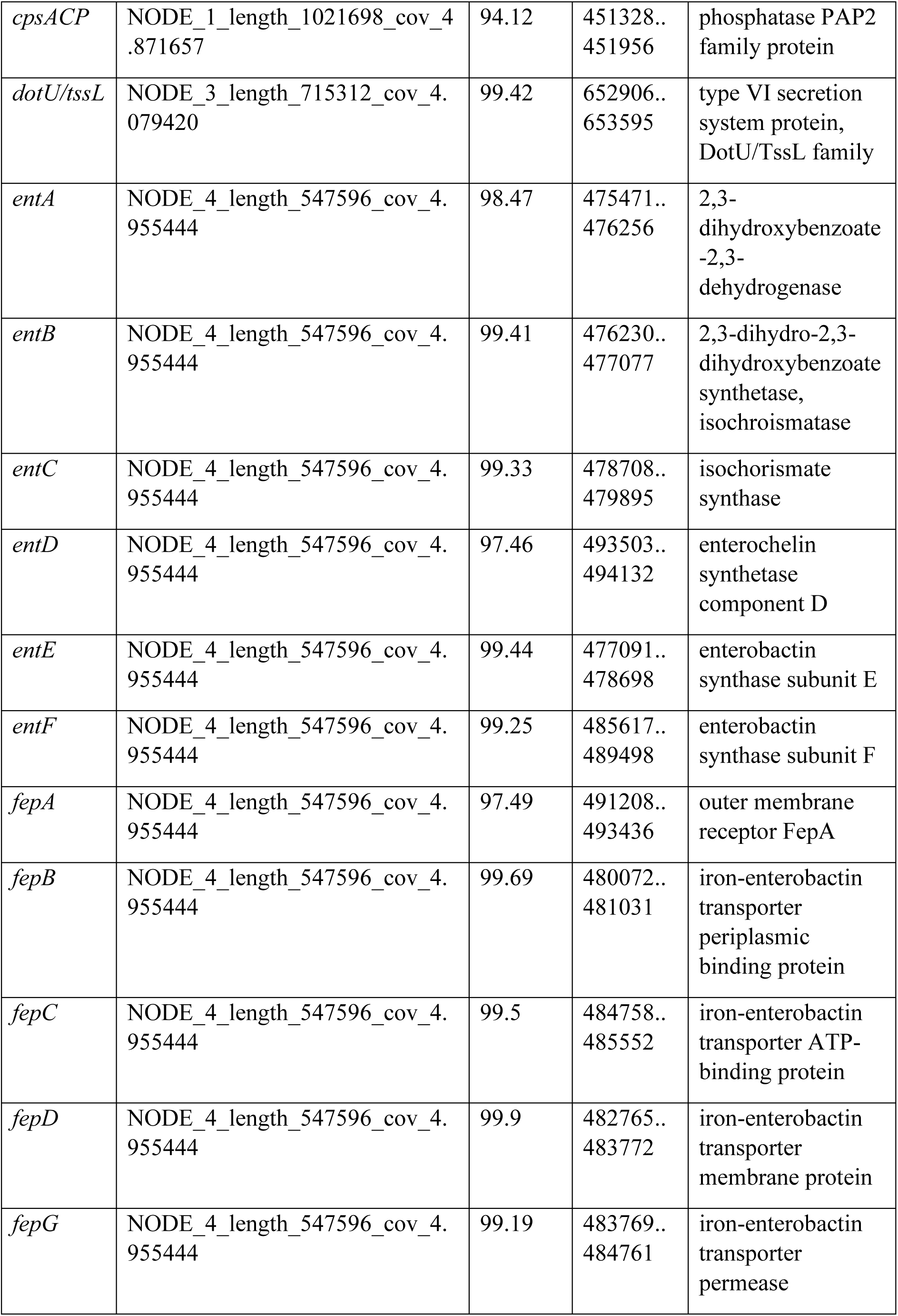
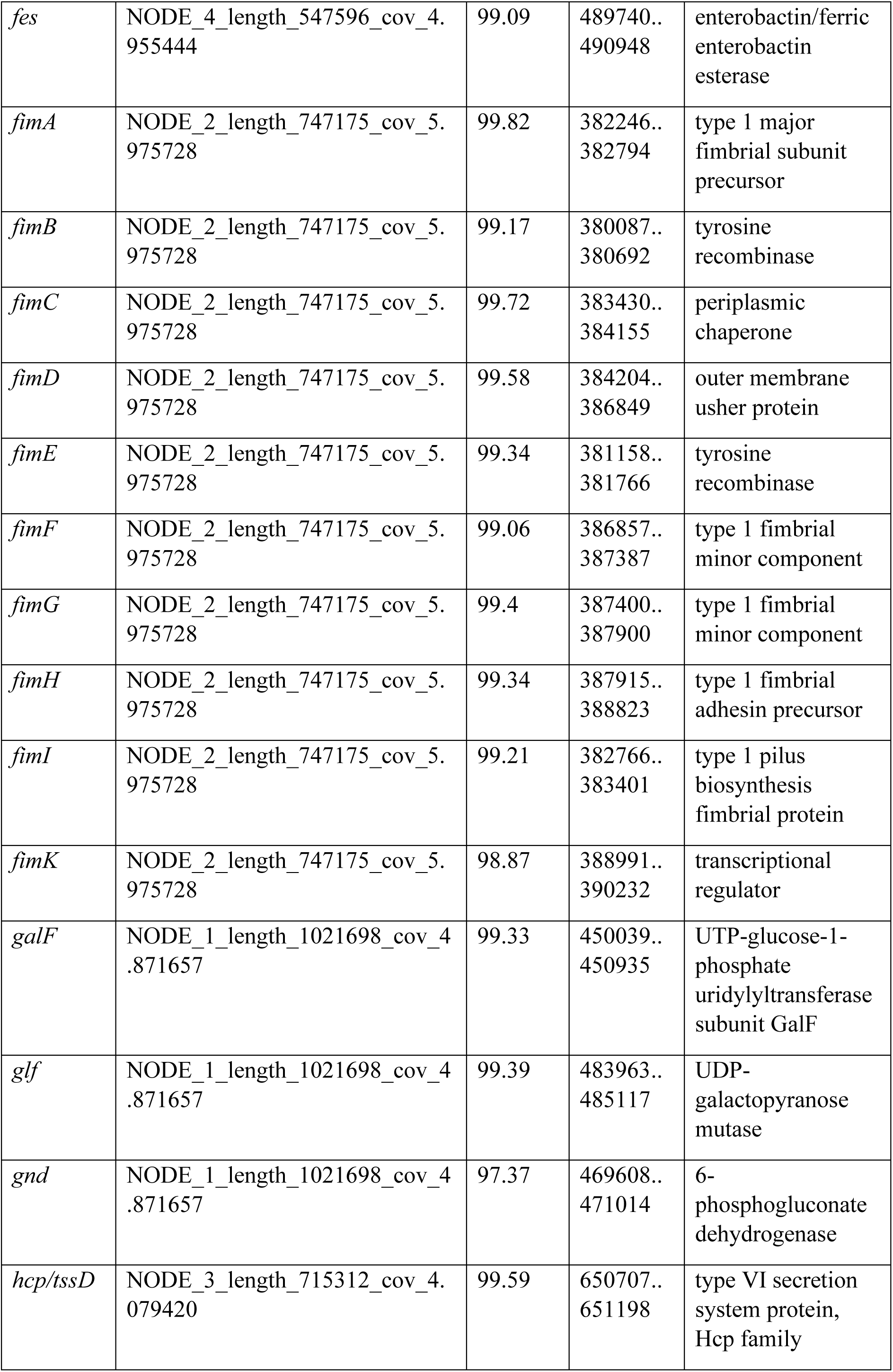
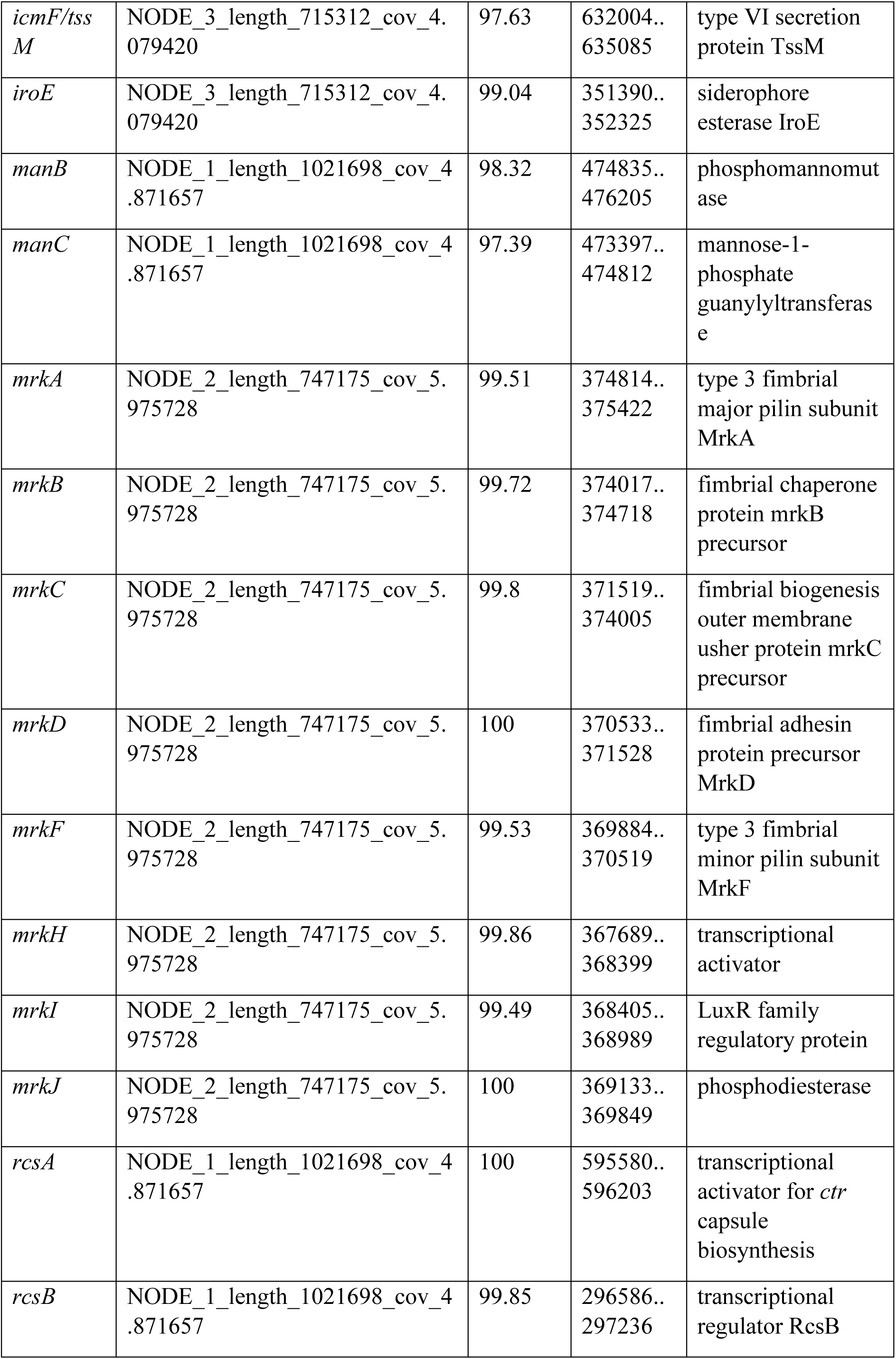
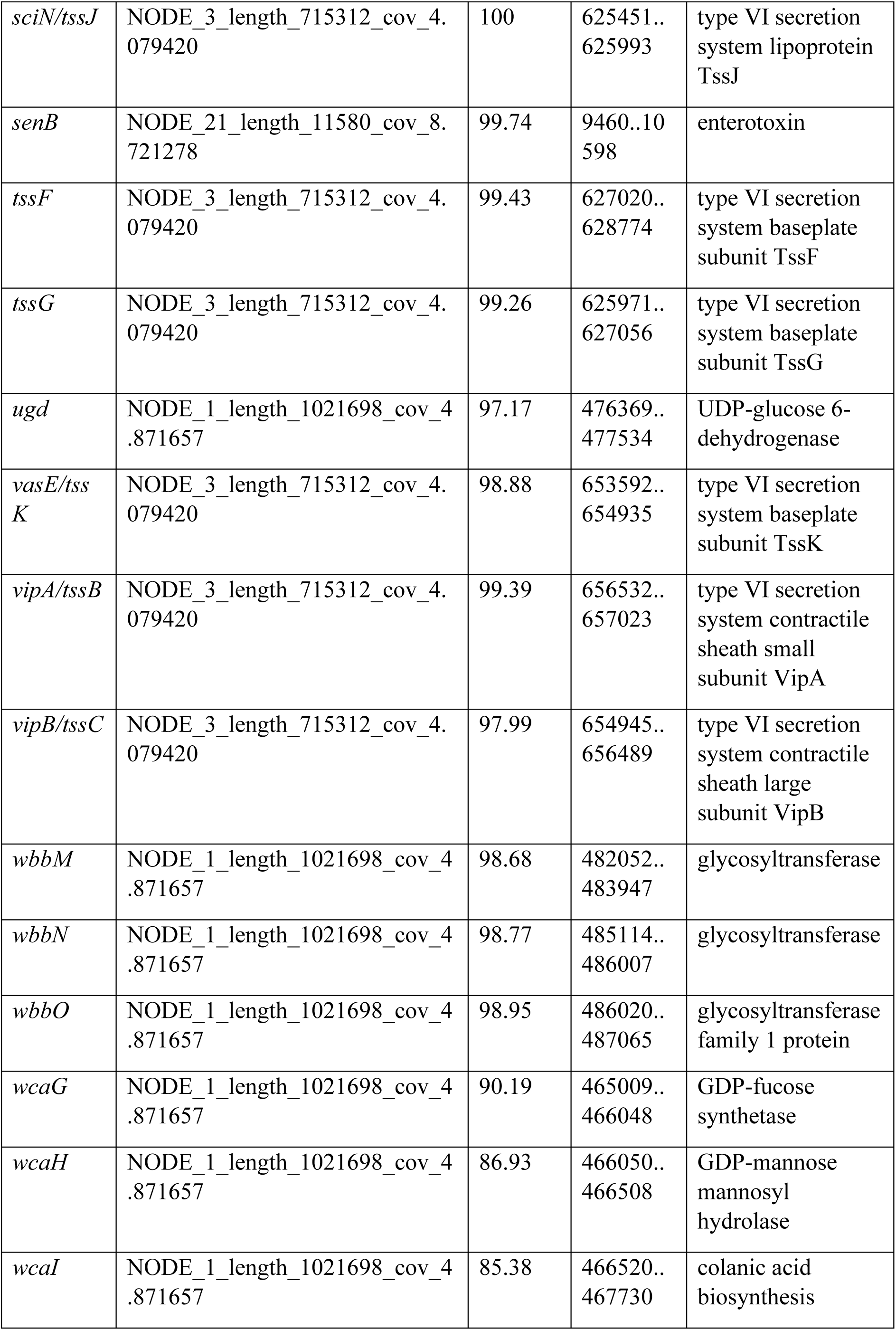
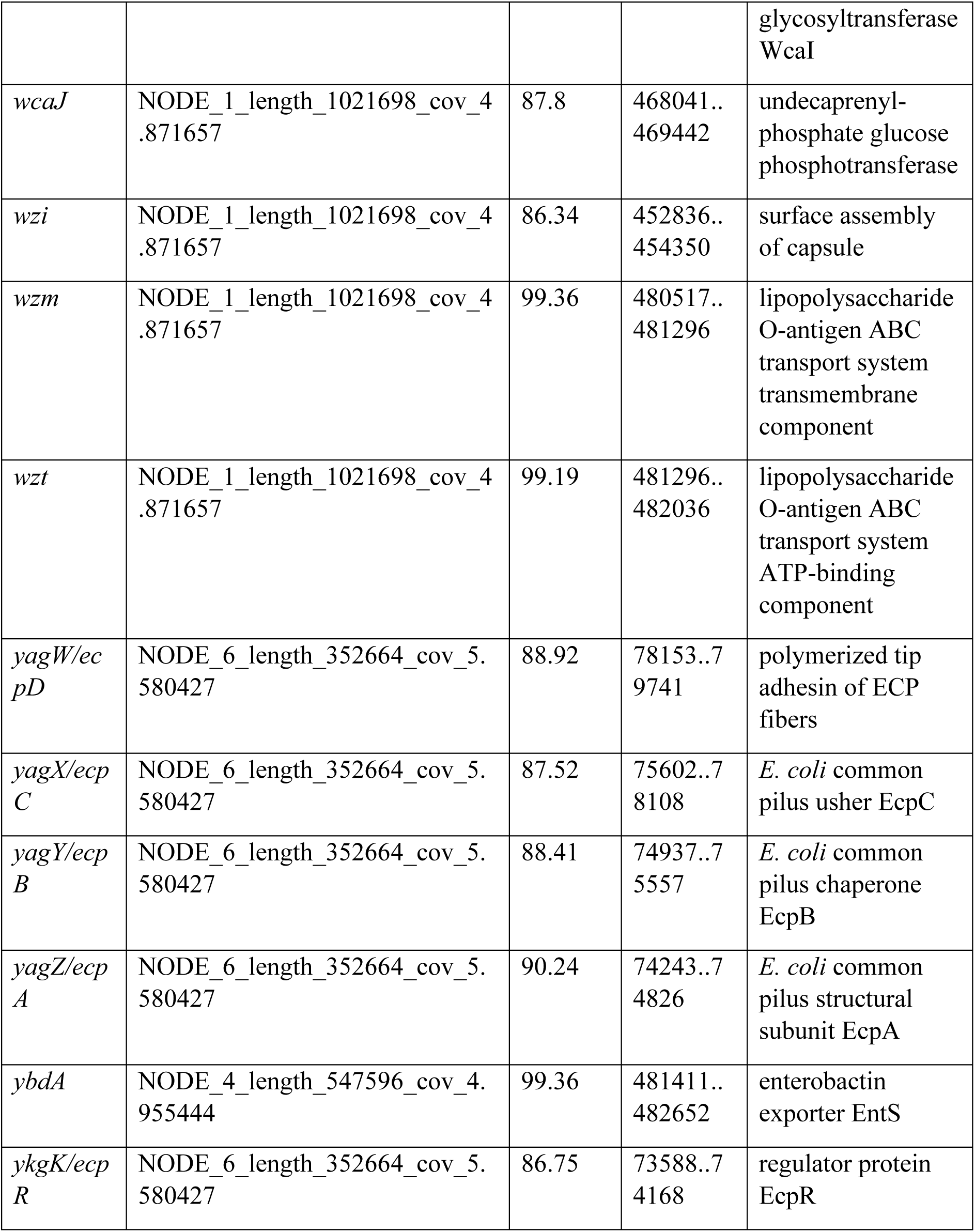
Identified Putative Virulence Genes in *K. pneumoniae* CP4. A total of 69 virulence genes were identified using the BacWGSTdb 2.0.

**Supplementary Table 5:**
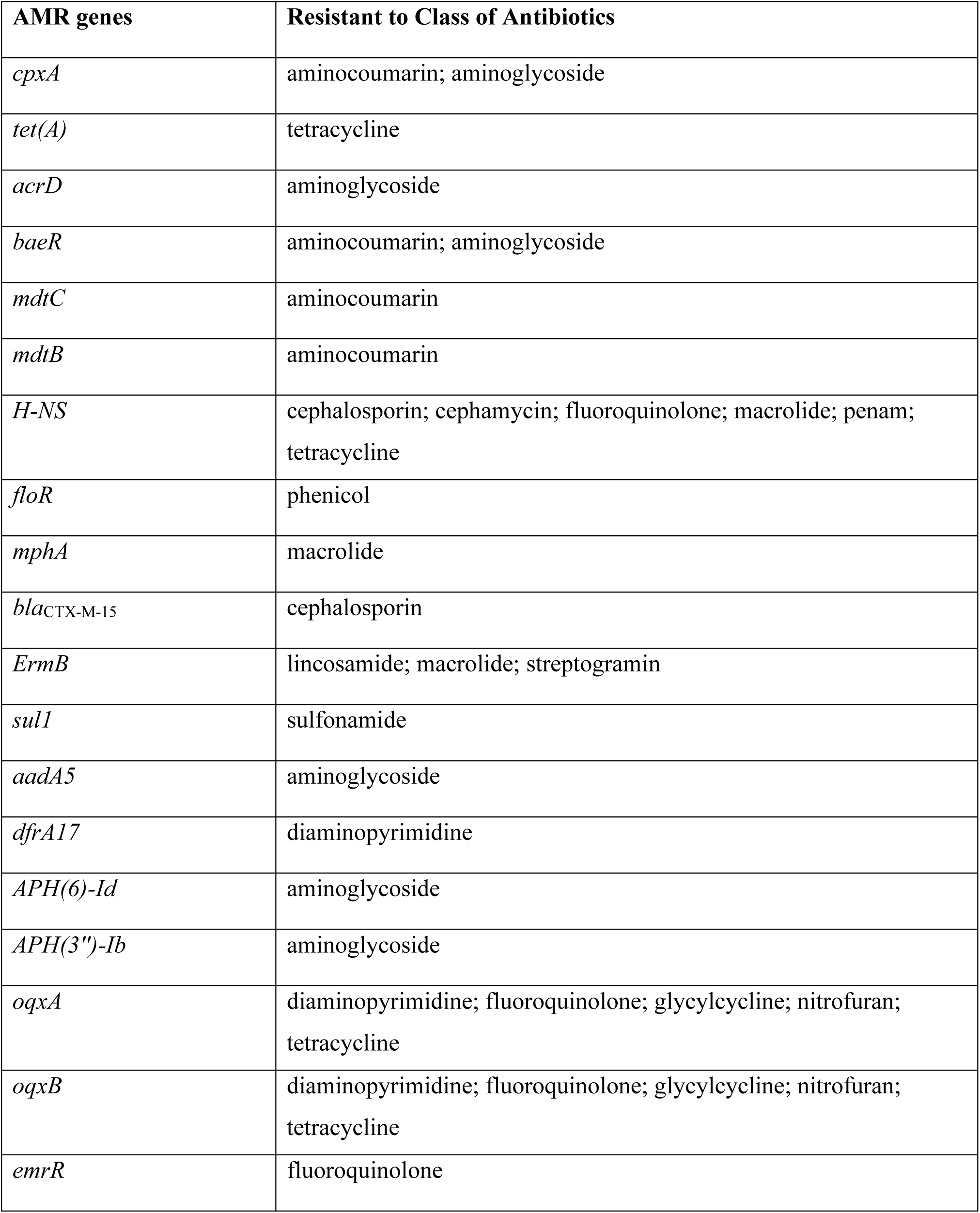
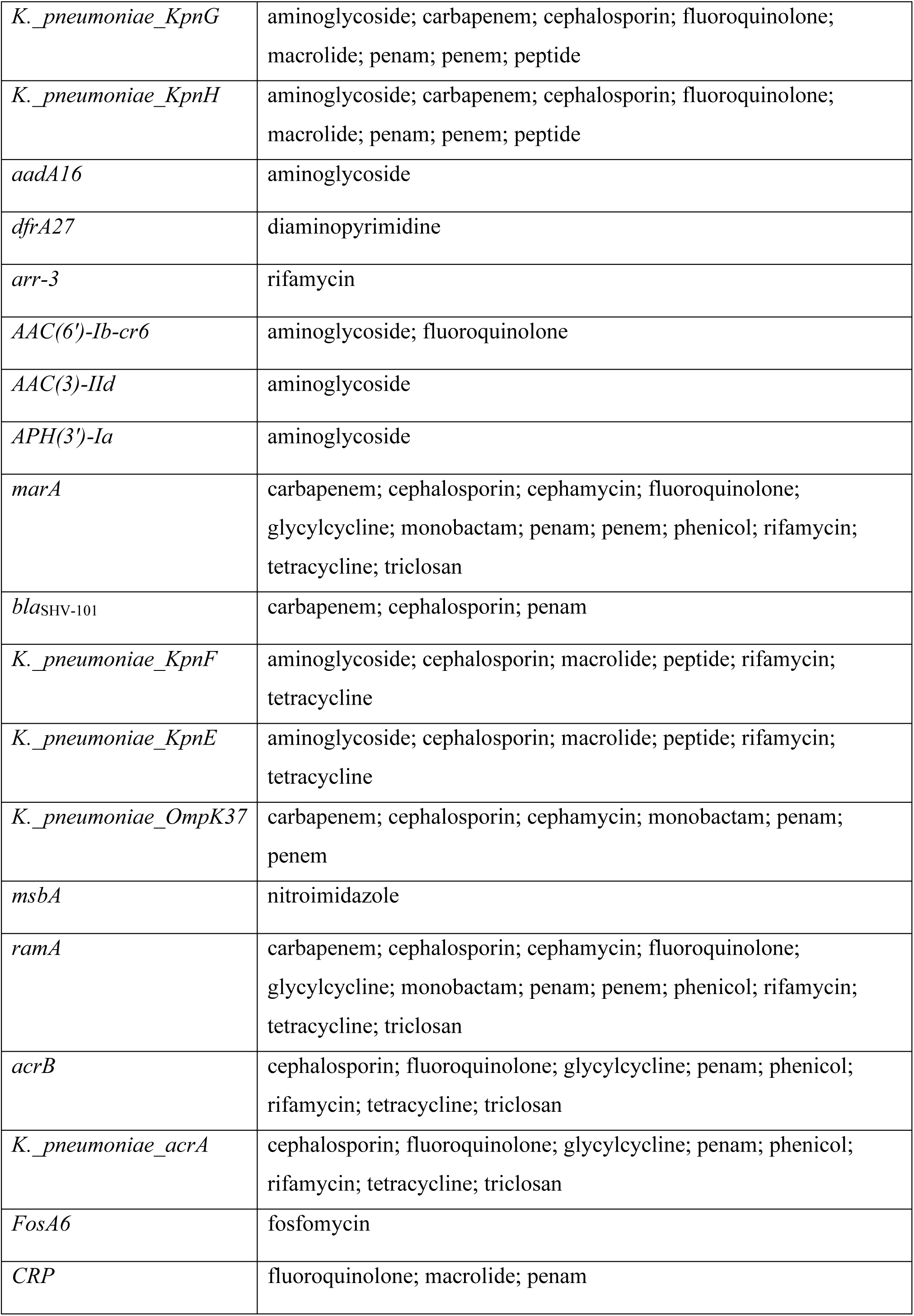
List of the putative Antimicrobial Resistance genes identified in the *K. pneumoniae* CP4 isolate using Abricate 1.0.1 against the CARD database, along with the corresponding classes of antibiotics to which each gene confers resistance.

**Supplementary Table 6:**
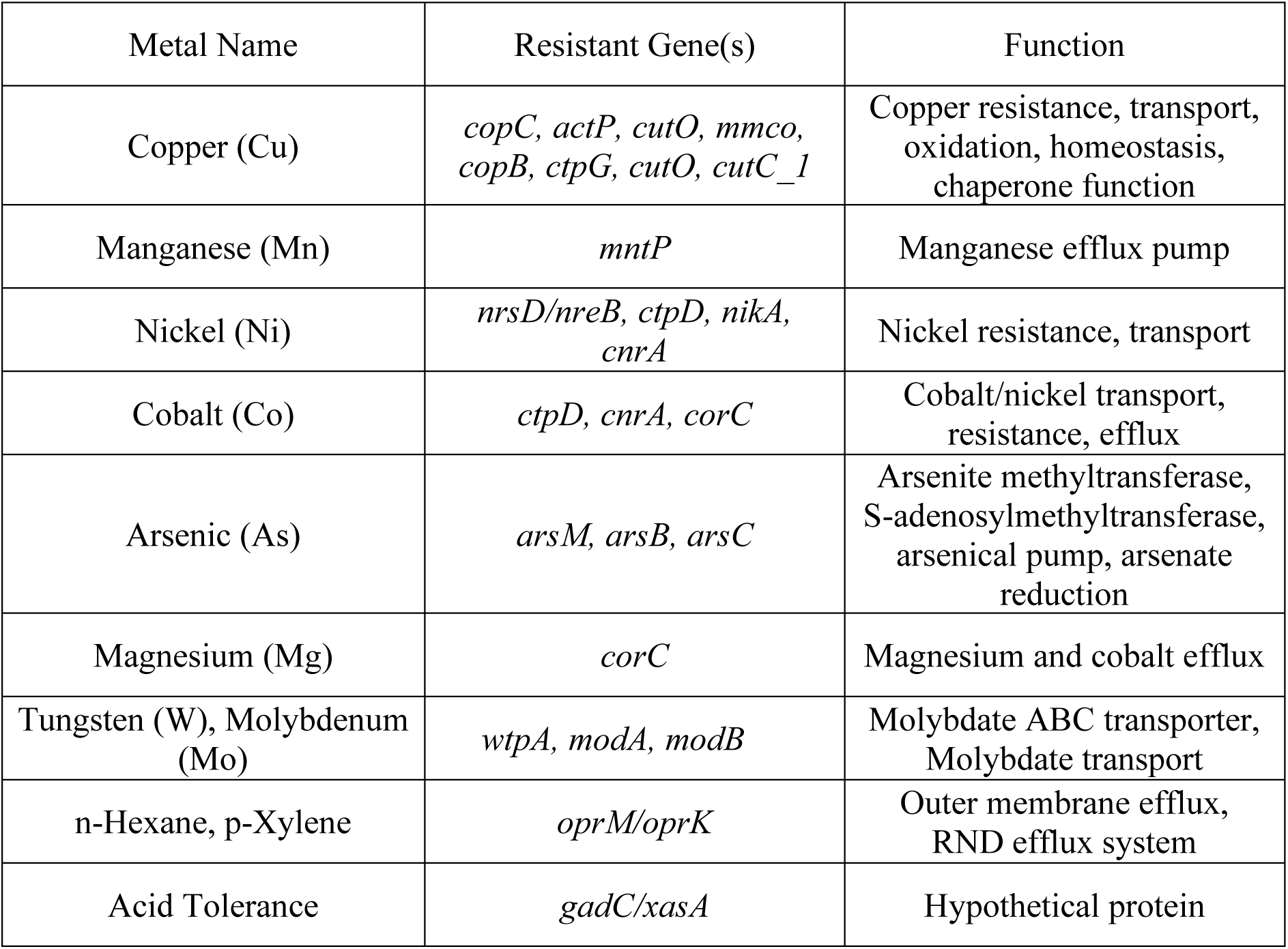
Identification of metal and putative chemical-resistant genes in *K. pneumoniae strain* CP4 was performed using BLASTP against the BacMet database. The BLOSUM80 substitution matrix was applied, and hits were filtered based on an E-value threshold of 0.01 to ensure significance. The identified genes and their corresponding functions related to metal and chemical resistance are presented.

**Supplementary Table 7:** Identified putative plasmid replicon sequence types in *Klebsiella pneumoniae CP4* genome by ABRicate 1.0.1 Using the PlasmidFinder Database.

| Putative plasmid replicon sequence types | %COVERAGE | %IDENTITY |
| --- | --- | --- |
| IncFIB(AP001918)_1 | 100 | 96.63 |
| IncFII(29)_1_pUTI89 | 100 | 99.61 |
| IncR_1 | 100 | 100 |
| Col156_1 | 92.21 | 98.59 |
| ColRNAI_1 | 98.46 | 81.39 |
| Col440II_1 | 99.29 | 83.75 |

**Supplementary Table 8:**
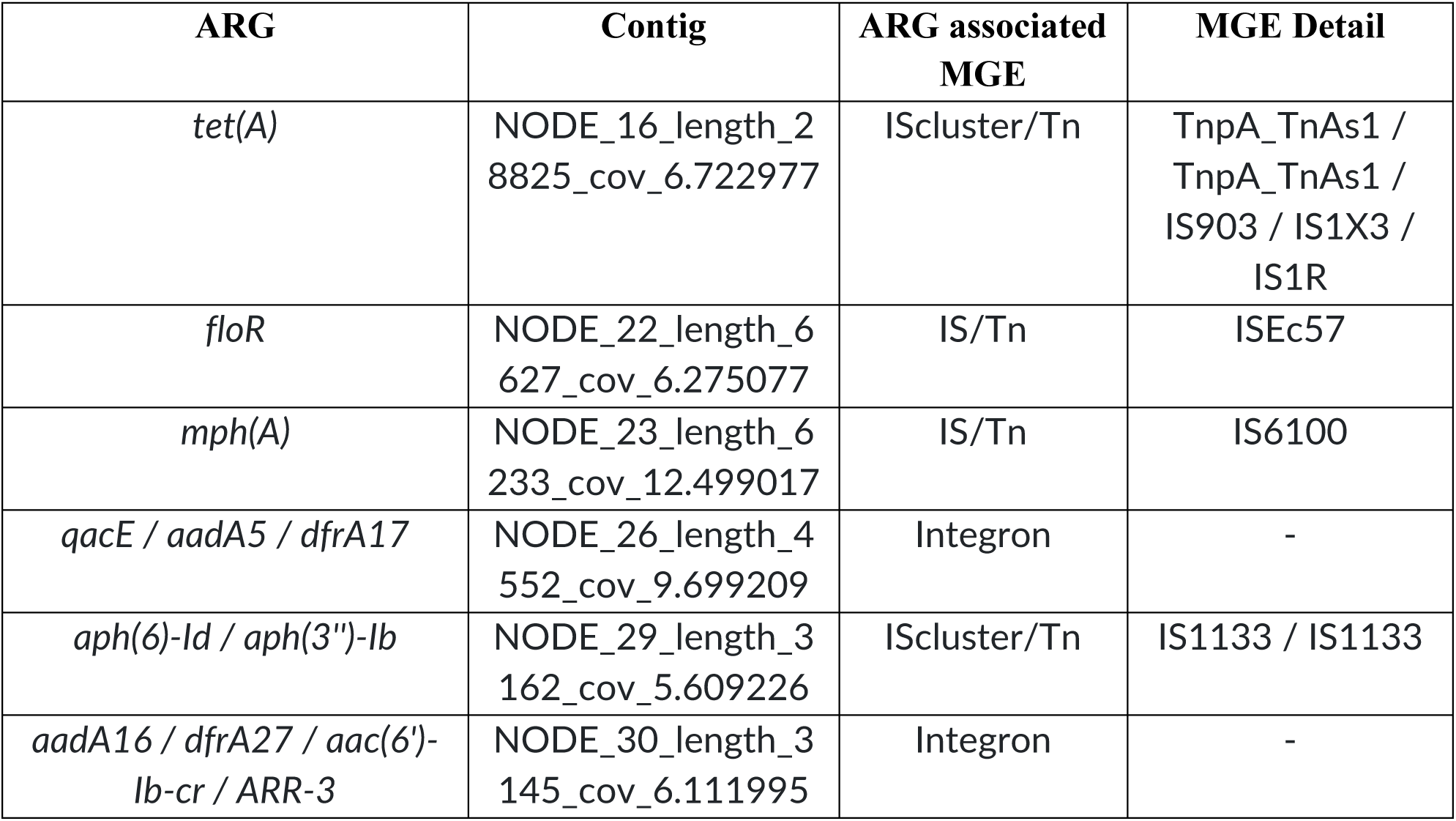
Putative Mobile Genetic Elements (MGE) of *K. pneumoniae* CP4 Carrying the Antibiotic Resistance Genes (ARG) Identified by VRprofile 2.

**Supplementary Table 9:**
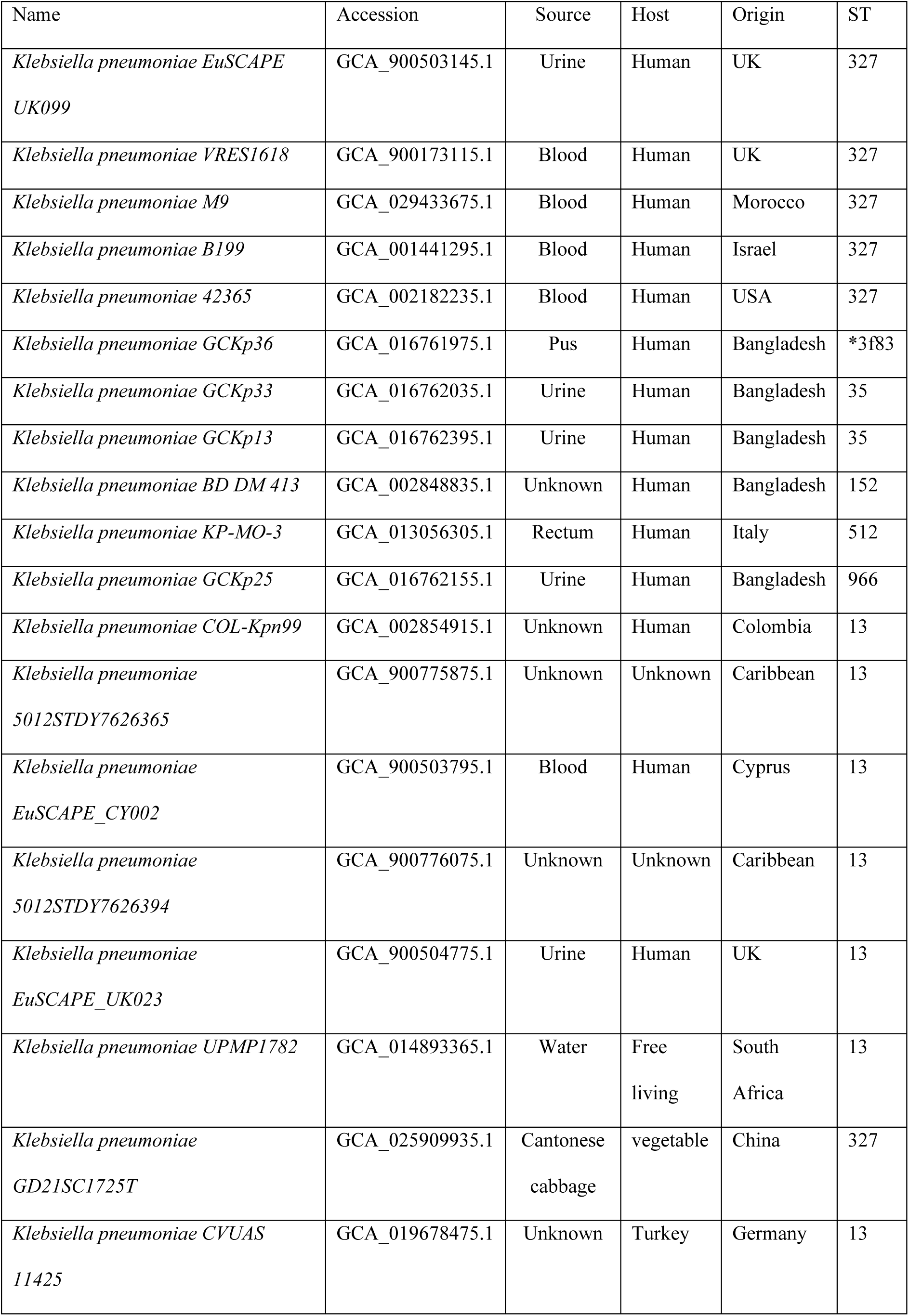
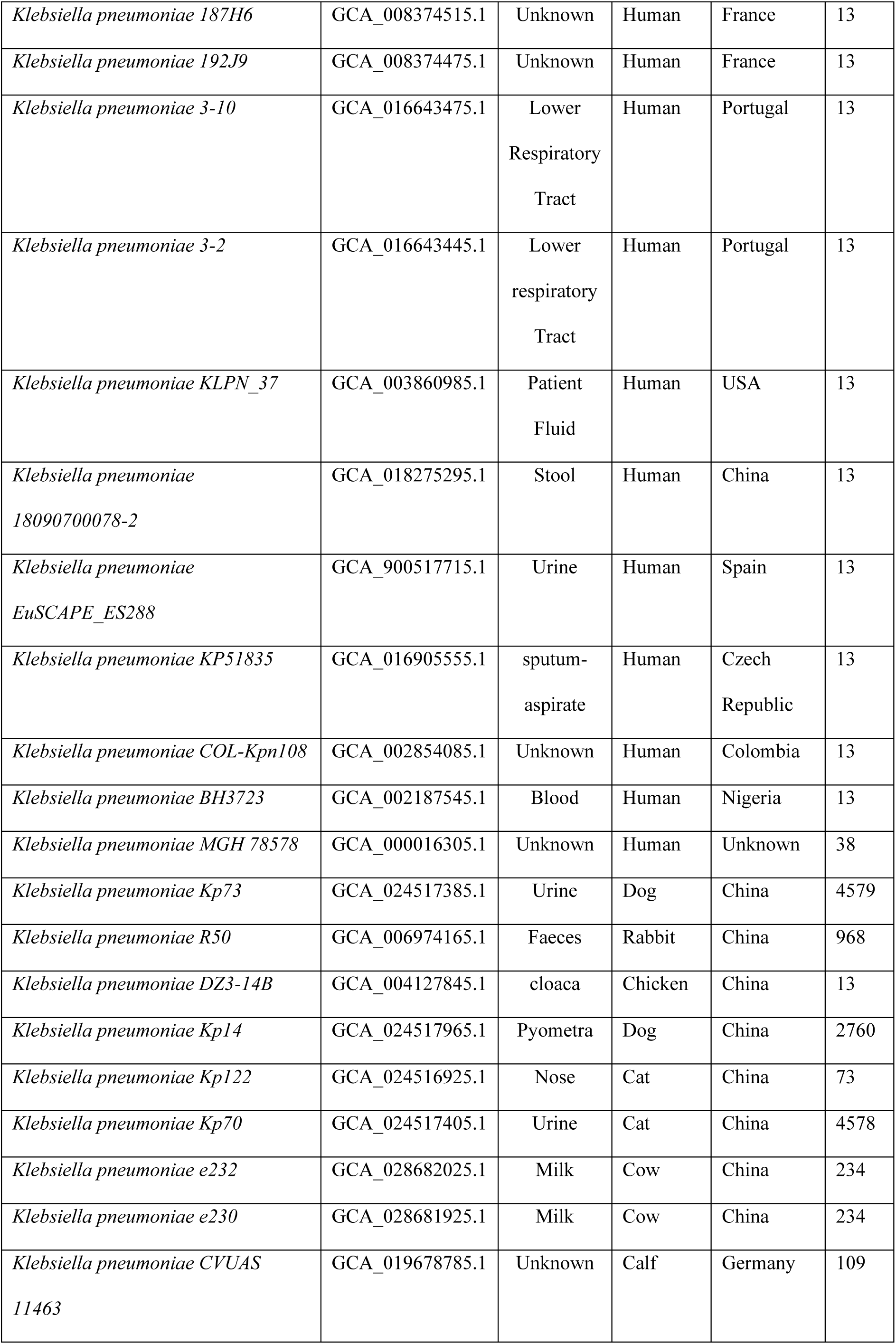
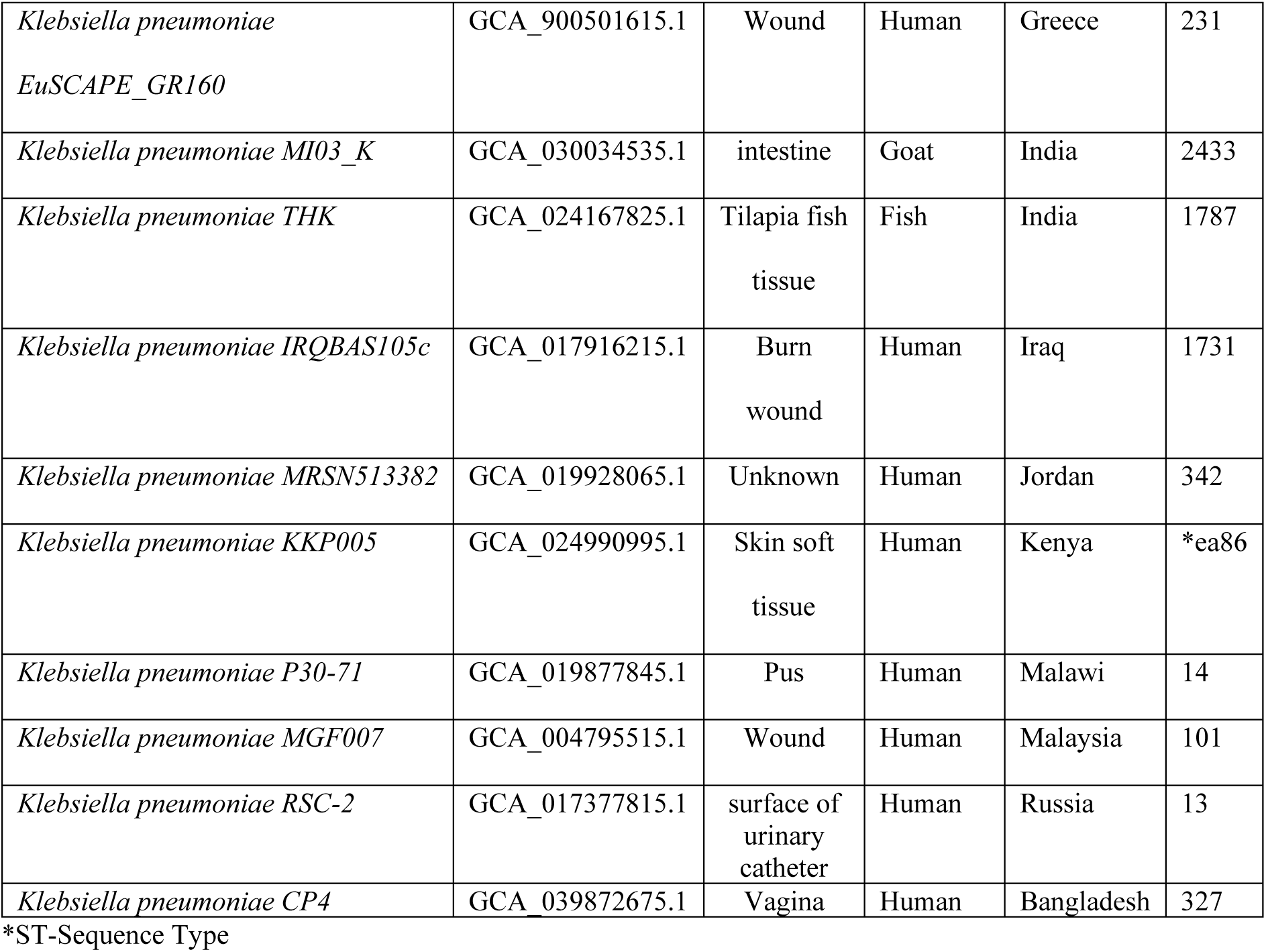
Sequences Used in Phylogenetic and Pangenome Analysis.

| Name | Accession | Source | Host | Origin | ST |
| --- | --- | --- | --- | --- | --- |
| *Klebsiella pneumoniae* EuSCAPE<br>UK099 | GCA_900503145.1 | Urine | Human | UK | 327 |
| *Klebsiella pneumoniae* VRES1618 | GCA_900173115.1 | Blood | Human | UK | 327 |
| *Klebsiella pneumoniae* M9 | GCA_029433675.1 | Blood | Human | Morocco | 327 |
| *Klebsiella pneumoniae* B199 | GCA_001441295.1 | Blood | Human | Israel | 327 |
| *Klebsiella pneumoniae* 42365 | GCA_002182235.1 | Blood | Human | USA | 327 |
| *Klebsiella pneumoniae* GCKp36 | GCA_016761975.1 | Pus | Human | Bangladesh | *3f83 |
| *Klebsiella pneumoniae* GCKp33 | GCA_016762035.1 | Urine | Human | Bangladesh | 35 |
| *Klebsiella pneumoniae* GCKp13 | GCA_016762395.1 | Urine | Human | Bangladesh | 35 |
| *Klebsiella pneumoniae* BD DM 413 | GCA_002848835.1 | Unknown | Human | Bangladesh | 152 |
| *Klebsiella pneumoniae* KP-MO-3 | GCA_013056305.1 | Rectum | Human | Italy | 512 |
| *Klebsiella pneumoniae* GCKp25 | GCA_016762155.1 | Urine | Human | Bangladesh | 966 |
| *Klebsiella pneumoniae* COL-Kpn99 | GCA_002854915.1 | Unknown | Human | Colombia | 13 |
| *Klebsiella pneumoniae*<br>5012STDY7626365 | GCA_900775875.1 | Unknown | Unknown | Caribbean | 13 |
| *Klebsiella pneumoniae*<br>EuSCAPE_CY002 | GCA_900503795.1 | Blood | Human | Cyprus | 13 |
| *Klebsiella pneumoniae*<br>5012STDY7626394 | GCA_900776075.1 | Unknown | Unknown | Caribbean | 13 |
| *Klebsiella pneumoniae*<br>EuSCAPE_UK023 | GCA_900504775.1 | Urine | Human | UK | 13 |
| *Klebsiella pneumoniae* UPMP1782 | GCA_014893365.1 | Water | Free<br>living | South<br>Africa | 13 |
| *Klebsiella pneumoniae*<br>GD21SC1725T | GCA_025909935.1 | Cantonese<br>cabbage | vegetable | China | 327 |
| *Klebsiella pneumoniae* CVUAS<br>11425 | GCA_019678475.1 | Unknown | Turkey | Germany | 13 |
| *Klebsiella pneumoniae* 187H6 | GCA_008374515.1 | Unknown | Human | France | 13 |
| *Klebsiella pneumoniae* 192J9 | GCA_008374475.1 | Unknown | Human | France | 13 |
| *Klebsiella pneumoniae* 3-10 | GCA_016643475.1 | Lower<br>Respiratory<br>Tract | Human | Portugal | 13 |
| *Klebsiella pneumoniae* 3-2 | GCA_016643445.1 | Lower<br>respiratory<br>Tract | Human | Portugal | 13 |
| *Klebsiella pneumoniae* KLPN_37 | GCA_003860985.1 | Patient<br>Fluid | Human | USA | 13 |
| *Klebsiella pneumoniae*<br>18090700078-2 | GCA_018275295.1 | Stool | Human | China | 13 |
| *Klebsiella pneumoniae*<br>EuSCAPE_ES288 | GCA_900517715.1 | Urine | Human | Spain | 13 |
| *Klebsiella pneumoniae* KP51835 | GCA_016905555.1 | sputum-<br>aspirate | Human | Czech<br>Republic | 13 |
| *Klebsiella pneumoniae* COL-Kpn108 | GCA_002854085.1 | Unknown | Human | Colombia | 13 |
| *Klebsiella pneumoniae* BH3723 | GCA_002187545.1 | Blood | Human | Nigeria | 13 |
| *Klebsiella pneumoniae* MGH 78578 | GCA_000016305.1 | Unknown | Human | Unknown | 38 |
| *Klebsiella pneumoniae* Kp73 | GCA_024517385.1 | Urine | Dog | China | 4579 |
| *Klebsiella pneumoniae* R50 | GCA_006974165.1 | Faeces | Rabbit | China | 968 |
| *Klebsiella pneumoniae* DZ3-14B | GCA_004127845.1 | cloaca | Chicken | China | 13 |
| *Klebsiella pneumoniae* Kp14 | GCA_024517965.1 | Pyometra | Dog | China | 2760 |
| *Klebsiella pneumoniae* Kp122 | GCA_024516925.1 | Nose | Cat | China | 73 |
| *Klebsiella pneumoniae* Kp70 | GCA_024517405.1 | Urine | Cat | China | 4578 |
| *Klebsiella pneumoniae* e232 | GCA_028682025.1 | Milk | Cow | China | 234 |
| *Klebsiella pneumoniae* e230 | GCA_028681925.1 | Milk | Cow | China | 234 |
| *Klebsiella pneumoniae* CVUAS<br>11463 | GCA_019678785.1 | Unknown | Calf | Germany | 109 |
| *Klebsiella pneumoniae*<br>*EuSCAPE_GR160* | GCA_900501615.1 | Wound | Human | Greece | 231 |
| *Klebsiella pneumoniae* MI03_K | GCA_030034535.1 | intestine | Goat | India | 2433 |
| *Klebsiella pneumoniae* THK | GCA_024167825.1 | Tilapia fish<br>tissue | Fish | India | 1787 |
| *Klebsiella pneumoniae* IRQBAS105c | GCA_017916215.1 | Burn<br>wound | Human | Iraq | 1731 |
| *Klebsiella pneumoniae* MRSN513382 | GCA_019928065.1 | Unknown | Human | Jordan | 342 |
| *Klebsiella pneumoniae* KKP005 | GCA_024990995.1 | Skin soft<br>tissue | Human | Kenya | *ea86 |
| *Klebsiella pneumoniae* P30-71 | GCA_019877845.1 | Pus | Human | Malawi | 14 |
| *Klebsiella pneumoniae* MGF007 | GCA_004795515.1 | Wound | Human | Malaysia | 101 |
| *Klebsiella pneumoniae* RSC-2 | GCA_017377815.1 | surface of<br>urinary<br>catheter | Human | Russia | 13 |
| *Klebsiella pneumoniae* CP4 | GCA_039872675.1 | Vagina | Human | Bangladesh | 327 |

\*ST-Sequence Type

**Supplementary Table 10:**
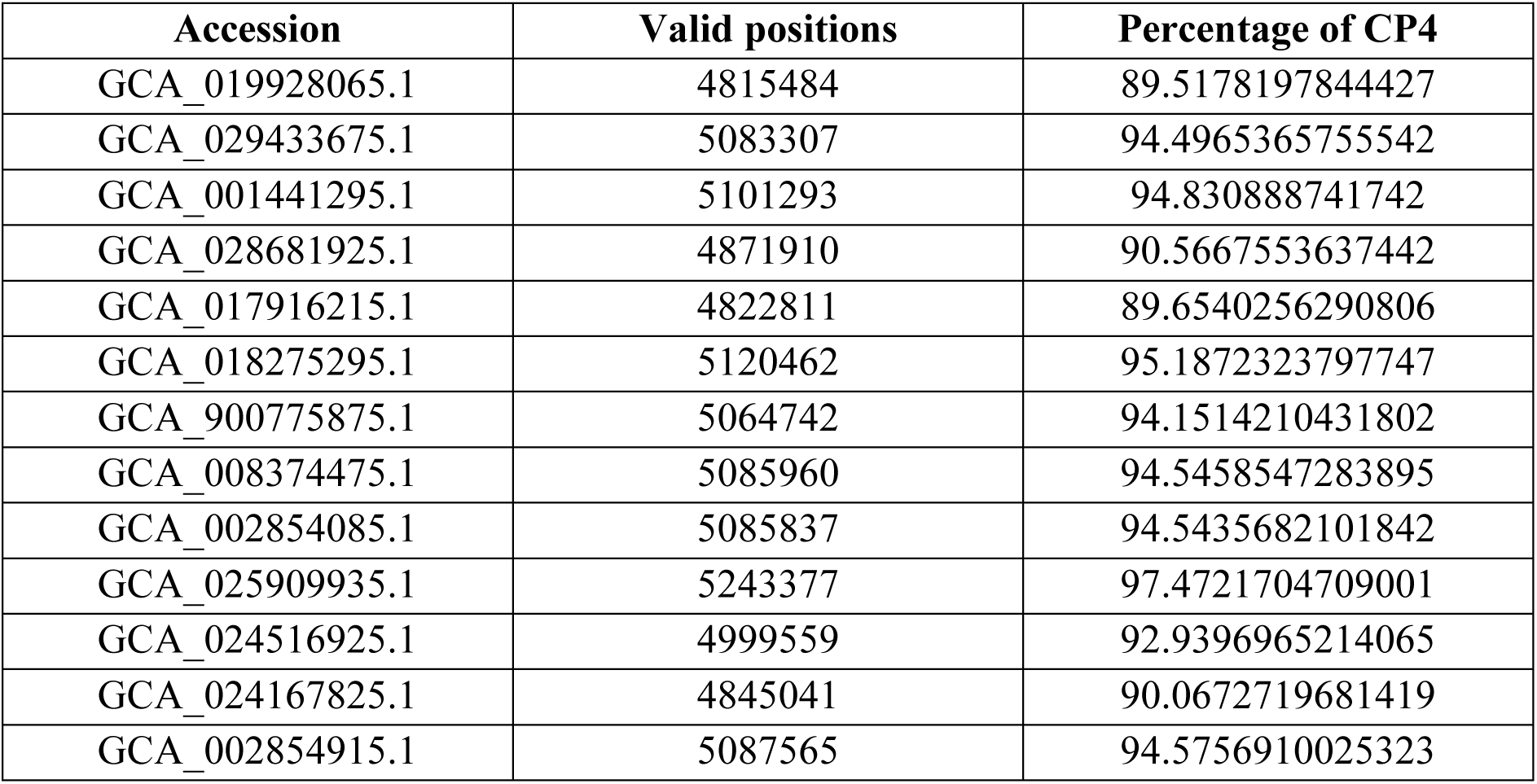
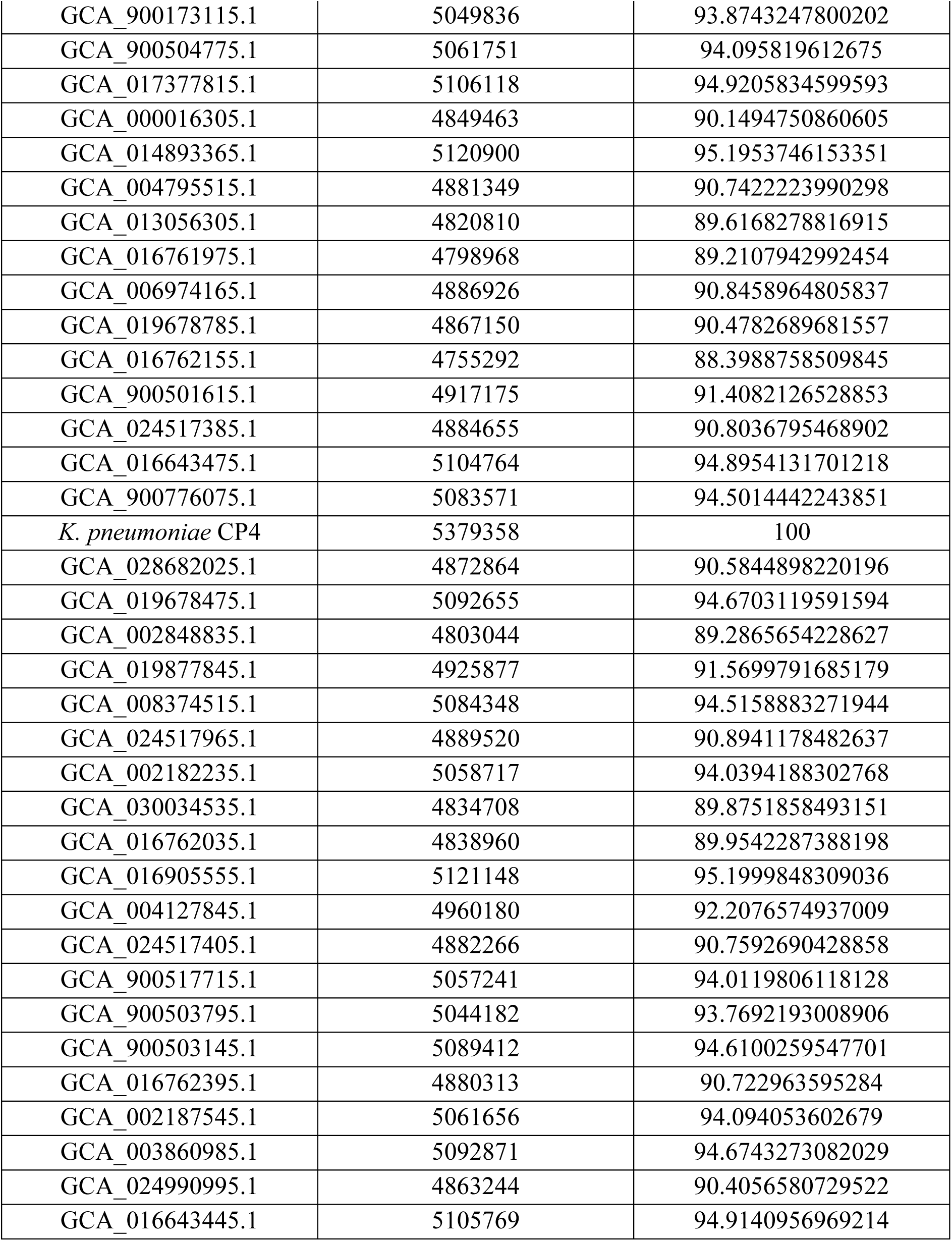
The number of positions that are shared and trusted between each isolate used in phylogenetic analysis and the *K. pneumoniae* CP4 genome.

| Accession | Valid positions | Percentage of CP4 |
| --- | --- | --- |
| GCA_019928065.1 | 4815484 | 89.5178197844427 |
| GCA_029433675.1 | 5083307 | 94.4965365755542 |
| GCA_001441295.1 | 5101293 | 94.830888741742 |
| GCA_028681925.1 | 4871910 | 90.5667553637442 |
| GCA_017916215.1 | 4822811 | 89.6540256290806 |
| GCA_018275295.1 | 5120462 | 95.1872323797747 |
| GCA_900775875.1 | 5064742 | 94.1514210431802 |
| GCA_008374475.1 | 5085960 | 94.5458547283895 |
| GCA_002854085.1 | 5085837 | 94.5435682101842 |
| GCA_025909935.1 | 5243377 | 97.4721704709001 |
| GCA_024516925.1 | 4999559 | 92.9396965214065 |
| GCA_024167825.1 | 4845041 | 90.0672719681419 |
| GCA_002854915.1 | 5087565 | 94.5756910025323 |
| GCA_900173115.1 | 5049836 | 93.8743247800202 |
| GCA_900504775.1 | 5061751 | 94.095819612675 |
| GCA_017377815.1 | 5106118 | 94.9205834599593 |
| GCA_000016305.1 | 4849463 | 90.1494750860605 |
| GCA_014893365.1 | 5120900 | 95.1953746153351 |
| GCA_004795515.1 | 4881349 | 90.7422223990298 |
| GCA_013056305.1 | 4820810 | 89.6168278816915 |
| GCA_016761975.1 | 4798968 | 89.2107942992454 |
| GCA_006974165.1 | 4886926 | 90.8458964805837 |
| GCA_019678785.1 | 4867150 | 90.4782689681557 |
| GCA_016762155.1 | 4755292 | 88.3988758509845 |
| GCA_900501615.1 | 4917175 | 91.4082126528853 |
| GCA_024517385.1 | 4884655 | 90.8036795468902 |
| GCA_016643475.1 | 5104764 | 94.8954131701218 |
| GCA_900776075.1 | 5083571 | 94.5014442243851 |
| *K. pneumoniae* CP4 | 5379358 | 100 |
| GCA_028682025.1 | 4872864 | 90.5844898220196 |
| GCA_019678475.1 | 5092655 | 94.6703119591594 |
| GCA_002848835.1 | 4803044 | 89.2865654228627 |
| GCA_019877845.1 | 4925877 | 91.5699791685179 |
| GCA_008374515.1 | 5084348 | 94.5158883271944 |
| GCA_024517965.1 | 4889520 | 90.8941178482637 |
| GCA_002182235.1 | 5058717 | 94.0394188302768 |
| GCA_030034535.1 | 4834708 | 89.8751858493151 |
| GCA_016762035.1 | 4838960 | 89.9542287388198 |
| GCA_016905555.1 | 5121148 | 95.1999848309036 |
| GCA_004127845.1 | 4960180 | 92.2076574937009 |
| GCA_024517405.1 | 4882266 | 90.7592690428858 |
| GCA_900517715.1 | 5057241 | 94.0119806118128 |
| GCA_900503795.1 | 5044182 | 93.7692193008906 |
| GCA_900503145.1 | 5089412 | 94.6100259547701 |
| GCA_016762395.1 | 4880313 | 90.722963595284 |
| GCA_002187545.1 | 5061656 | 94.094053602679 |
| GCA_003860985.1 | 5092871 | 94.6743273082029 |
| GCA_024990995.1 | 4863244 | 90.4056580729522 |
| GCA_016643445.1 | 5105769 | 94.9140956969214 |

## Supplementary Figures

**Supplementary Figure 1:**
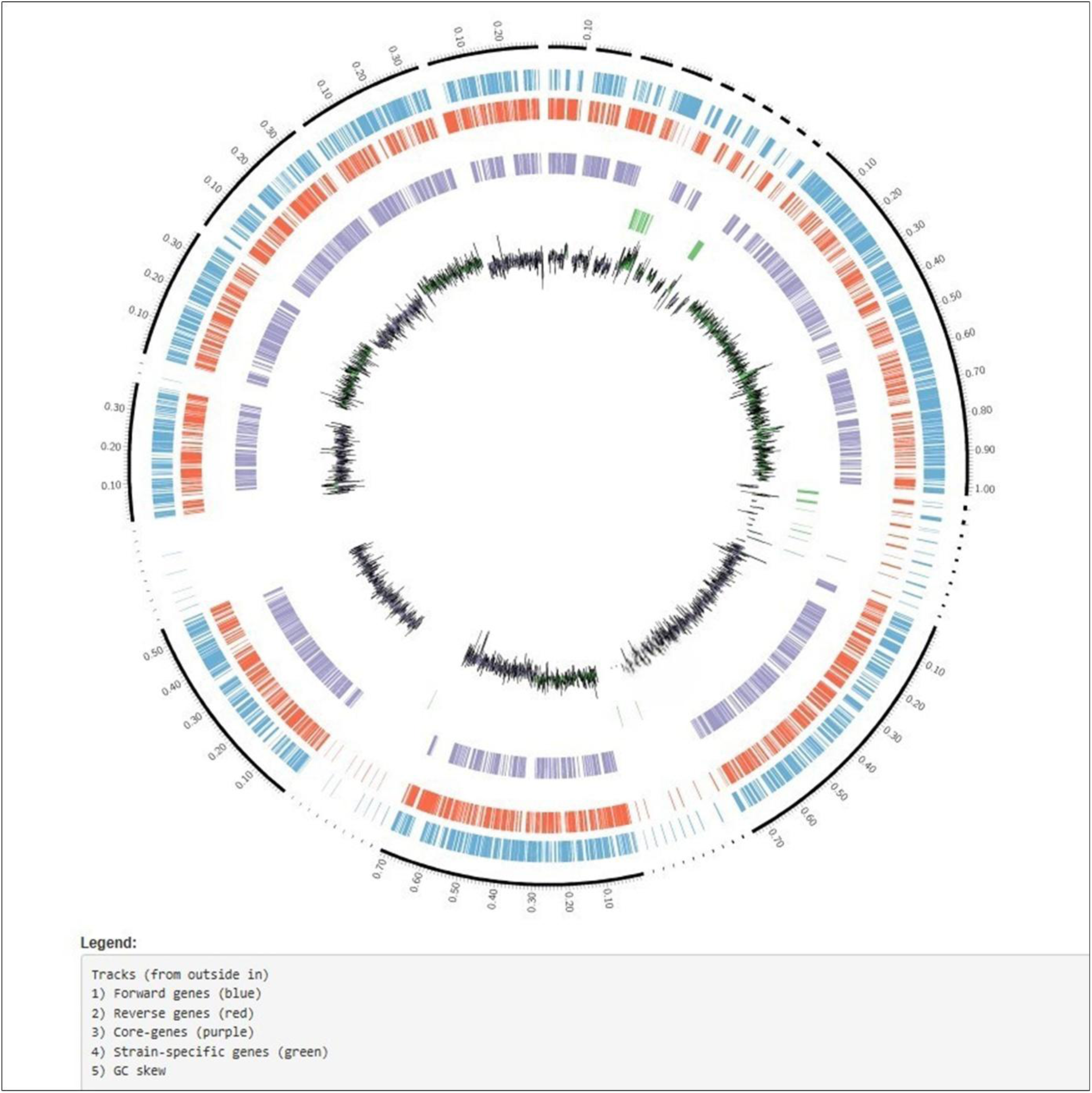
Circos plot generated by PanExplorer showing the genomic arrangement of *K. pneumoniae* CP4, including forward genes (blue), reverse genes (red), core genes (purple), strain-specific genes (green), and GC skew.

**Supplementary Figure 2:**
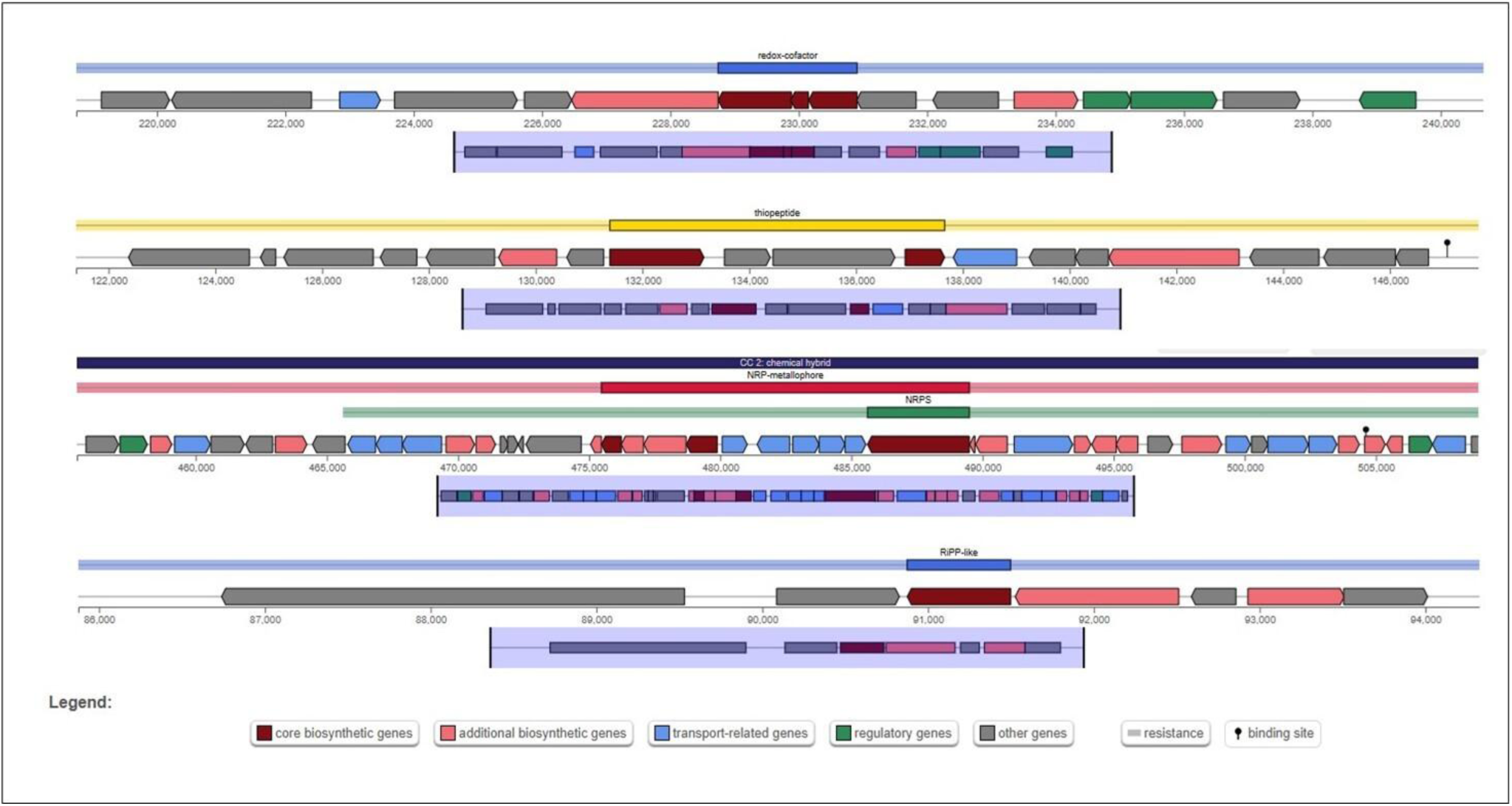
Identified putative secondary metabolite biosynthesis gene clusters including redox-cofactor, thiopeptide, NRP-metallophore, and RiPP-like clusters using antiSMASH bacterial version.

**Supplementary Figure 3:**
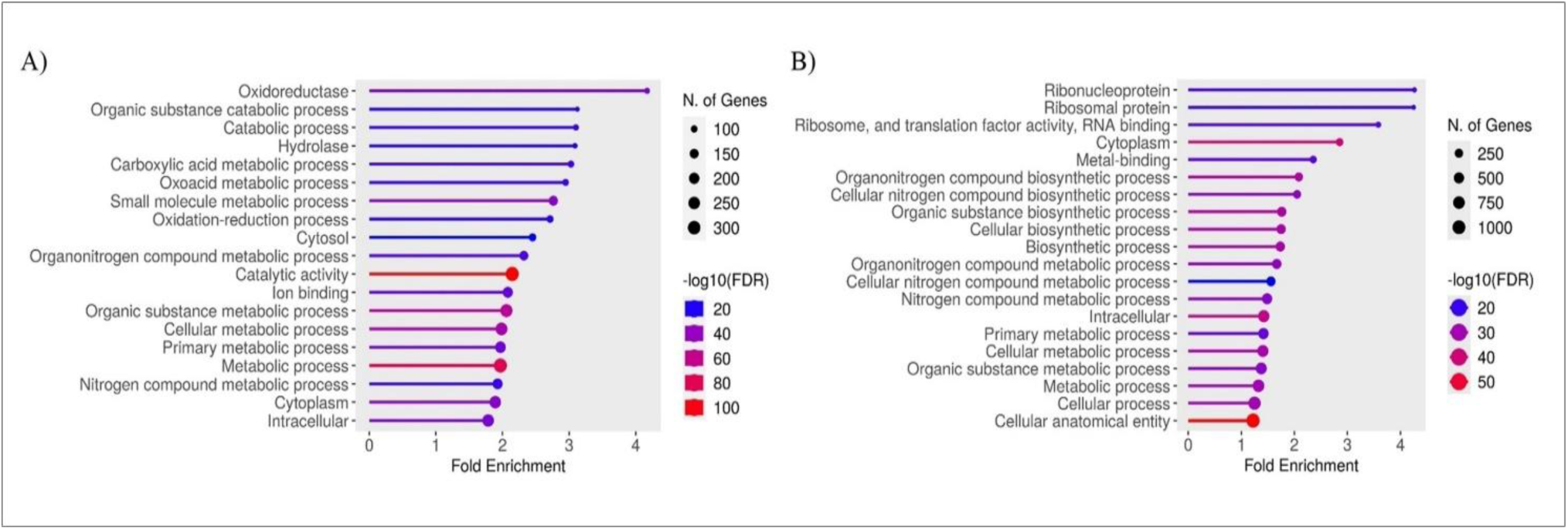
Pathway enrichment analysis performed using ShinyGO 0.80. A) Results based on EC numbers. B) Results based on gene-preferred names, annotated by eggNOG-mapper v2.

**Supplementary Figure 4:**
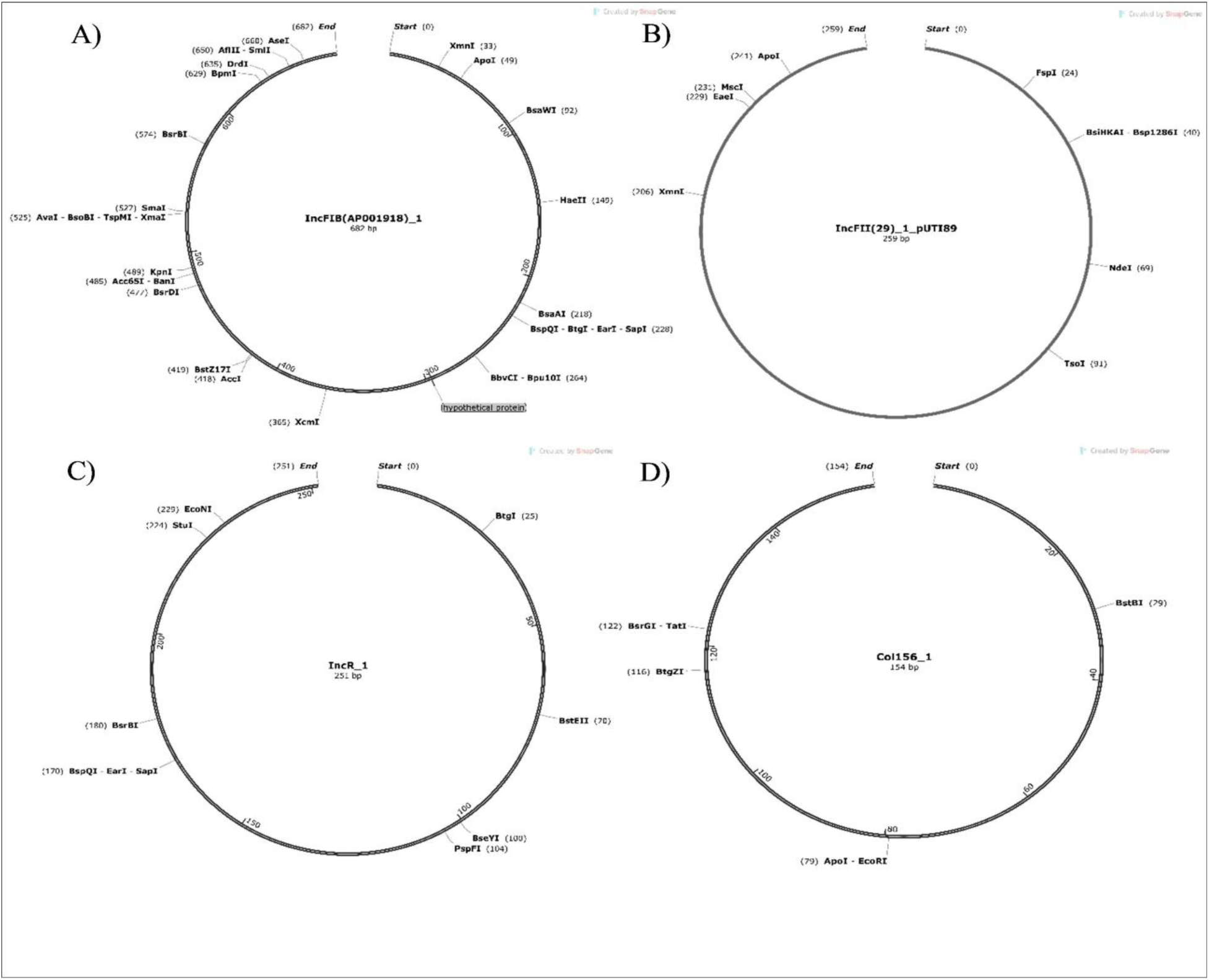
Visualization of selected putative plasmid replicon sequence types (>95% identity) identified in *Klebsiella pneumoniae* CP4, generated using SnapGene Viewer v7.2.1. Restriction enzyme recognition sites are indicated. **A)** IncFIB(AP001918)_1 **B)** IncFII(29)_1_pUTI89 **C)** IncR_1 **D)** Col156_1

**Supplementary Figure 5:**
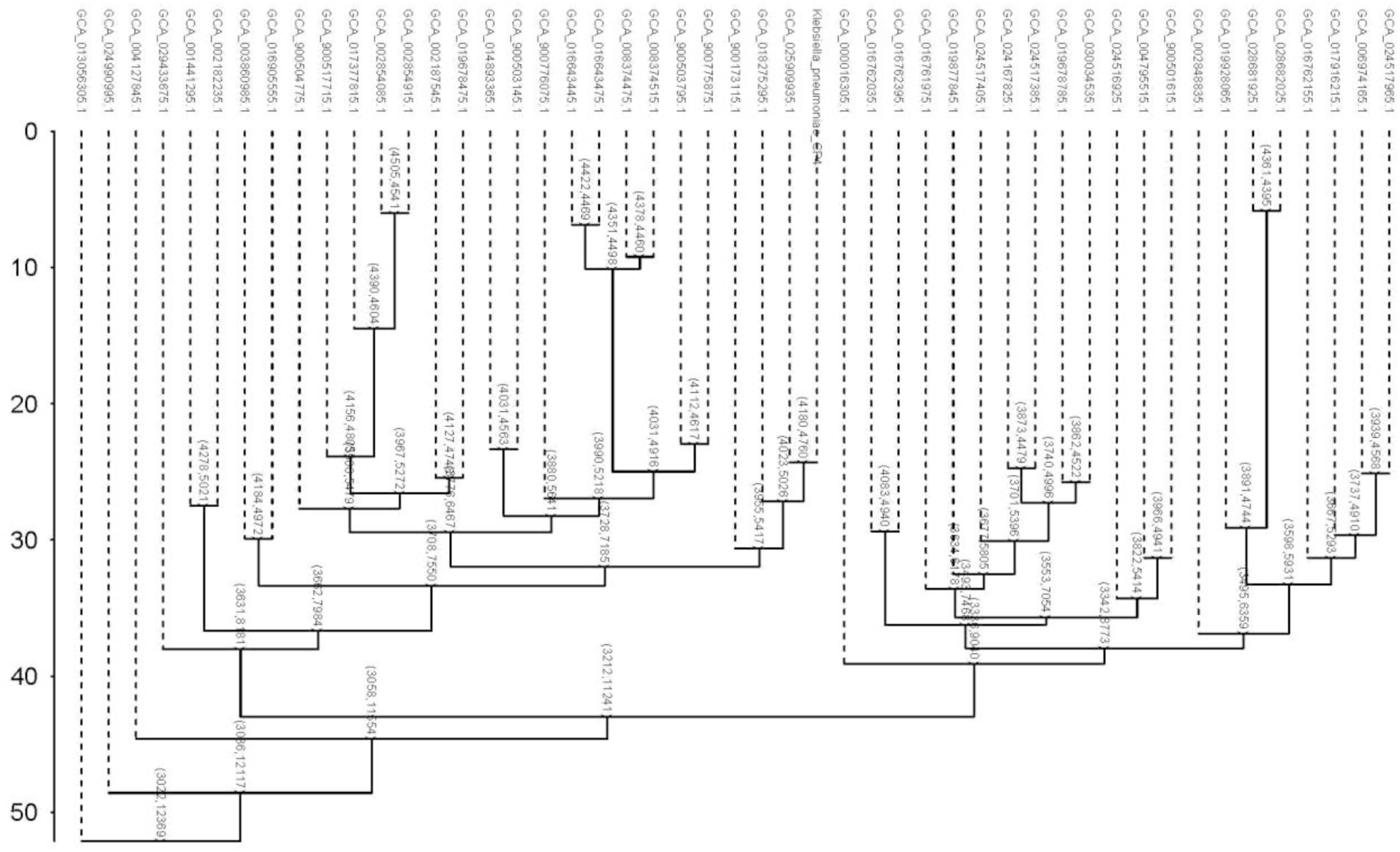
Pangenome analysis of *K. pneumoniae* using IPGA v1.09: Cluster sharing dendrogram showing the relationships among genomes based on shared gene clusters.

## Notes

https://www.ncbi.nlm.nih.gov/nuccore/JBDPJK000000000

### Competing Interest Statement

The authors have declared no competing interest.

### Summary of Updates

This version of the manuscript has been revised to update the images

### Funding

- University Grants Commission of Bangladesh: BIO-34

## References

[1] Piperaki E-T, Syrogiannopoulos GA, Tzouvelekis LS, Daikos GL. Klebsiella pneumoniae: Virulence, Biofilm and Antimicrobial Resistance. Pediatr Infect Dis J 2017;36.

[2] Mancuso G, Midiri A, Gerace E, Biondo C. Bacterial Antibiotic Resistance: The Most Critical Pathogens. Pathogens 2021;10:1310. 10.3390/pathogens10101310.

[3] Gorrie CL, Mirčeta M, Wick RR, Judd LM, Lam MMC, Gomi R, et al. Genomic dissection of Klebsiella pneumoniae infections in hospital patients reveals insights into an opportunistic pathogen. Nat Commun 2022;13:3017. 10.1038/s41467-022-30717-6.

[4] Eger E, Schwabe M, Schulig L, Hübner N-O, Bohnert JA, Bornscheuer UT, et al. Extensively Drug-Resistant Klebsiella pneumoniae Counteracts Fitness and Virulence Costs That Accompanied Ceftazidime-Avibactam Resistance Acquisition. Microbiol Spectr 2022;10. 10.1128/spectrum.00148-22.

[5] Munita JM, Arias CA. Mechanisms of Antibiotic Resistance. Microbiol Spectr 2016;4. 10.1128/microbiolspec.VMBF-0016-2015.

[6] Mahmud ZH, Uddin SZ, Moniruzzaman M, Ali S, Hossain M, Islam MdT, et al. Healthcare Facilities as Potential Reservoirs of Antimicrobial Resistant Klebsiella pneumoniae: An Emerging Concern to Public Health in Bangladesh. Pharmaceuticals 2022;15:1116. 10.3390/ph15091116.

[7] Zuleta-González MC, Zapata-Salazar ME, Guerrero-Hurtado LS, Puerta-Suárez J, Cardona-Maya WD. Klebsiella pneumoniae and Streptococcus agalactiae: Passengers in the sperm travel. Arch Esp Urol 2019;72:939–47.

[8] Eze EM;, Unegbu VN;, Ezebialu CU;, Nneji IR. Prevalence of microorganisms associated with Pelvic inflammatory disease in reproductive aged women in Onitsha North, Anambra state, Nigeria. Novel Research in Microbiology Journal 2018;2:147–55. 10.21608/nrmj.2018.22707.

[9] Magdum M, Chowdhury MdAT, Begum N, Riya S. Types of Infertility and Its Risk Factors among Infertile Women: A Prospective Study in Dhaka City. J Biosci Med (Irvine) 2022;10:158–68. 10.4236/jbm.2022.104014.

[10] Hasan Z, Begum N, Ahmed S, Yasmin M. Association of opportunistic bacterial pathogens with female infertility: A case–control study. Journal of Obstetrics and Gynaecology Research 2025;51. 10.1111/jog.16243.

[11] Kar S, Kawser Z, Sridhar S, Mukta SA, Hasan N, Siddik AB, et al. High Prevalence of Carbapenem-resistant Klebsiella Pneumoniae in Fecal and Water Samples in Dhaka, Bangladesh. Open Forum Infect Dis 2024;11:ofae612. 10.1093/ofid/ofae612.

[12] Hasan Z, Netherland M, Hasan NA, Begum N, Yasmin M, Ahmed S. An insight into the vaginal microbiome of infertile women in Bangladesh using metagenomic approach. Front Cell Infect Microbiol 2024;14. 10.3389/fcimb.2024.1390088.

[13] Ahmed OB*, Dablool AS. Quality Improvement of the DNA extracted by boiling method in Gram negative bacteria. Int J Bioassays 2017;6:5347. 10.21746/ijbio.2017.04.004.

[14] Hou Q, Bai X, Li W, Gao X, Zhang F, Sun Z, et al. Design of Primers for Evaluation of Lactic Acid Bacteria Populations in Complex Biological Samples. Front Microbiol 2018;9. 10.3389/fmicb.2018.02045.

[15] Bauer AW, Kirby WMM, Sherris JC, Turck M. Antibiotic Susceptibility Testing by a Standardized Single Disk Method. Am J Clin Pathol 1966;45:493–6. 10.1093/ajcp/45.4_ts.493.

[16] Clinical and Laboratory Standards Institute (CLSI). Performance Standards for Antimicrobial Susceptibility Testing. 34th ed. CLSI supplement M100 (ISBN 978-1- 68440-220-5 [Print]; ISBN 978-1-68440-221-2 [Electronic]). USA: Clinical and Laboratory Standards Institute; 2024 n.d.

[17] Chen S, Zhou Y, Chen Y, Gu J. fastp: an ultra-fast all-in-one FASTQ preprocessor. Bioinformatics 2018;34:i884–90. 10.1093/bioinformatics/bty560.

[18] Bankevich A, Nurk S, Antipov D, Gurevich AA, Dvorkin M, Kulikov AS, et al. SPAdes: A New Genome Assembly Algorithm and Its Applications to Single-Cell Sequencing. Journal of Computational Biology 2012;19:455–77. 10.1089/cmb.2012.0021.

[19] Mikheenko A, Prjibelski A, Saveliev V, Antipov D, Gurevich A. Versatile genome assembly evaluation with QUAST-LG. Bioinformatics 2018;34:i142–50. 10.1093/bioinformatics/bty266.

[20] Parks DH, Imelfort M, Skennerton CT, Hugenholtz P, Tyson GW. CheckM: assessing the quality of microbial genomes recovered from isolates, single cells, and metagenomes. Genome Res 2015;25:1043–55. 10.1101/gr.186072.114.

[21] Manni M, Berkeley MR, Seppey M, Simão FA, Zdobnov EM. BUSCO Update: Novel and Streamlined Workflows along with Broader and Deeper Phylogenetic Coverage for Scoring of Eukaryotic, Prokaryotic, and Viral Genomes. Mol Biol Evol 2021;38:4647–54. 10.1093/molbev/msab199.

[22] Seemann T. Prokka: rapid prokaryotic genome annotation. Bioinformatics 2014;30:2068–9. 10.1093/bioinformatics/btu153.

[23] Grant JR, Enns E, Marinier E, Mandal A, Herman EK, Chen C, et al. Proksee: in-depth characterization and visualization of bacterial genomes. Nucleic Acids Res 2023;51:W484–92. 10.1093/nar/gkad326.

[24] Aziz RK, Bartels D, Best AA, DeJongh M, Disz T, Edwards RA, et al. The RAST Server: Rapid Annotations using Subsystems Technology. BMC Genomics 2008;9:75. 10.1186/1471-2164-9-75.

[25] Dereeper A, Summo M, Meyer DF. PanExplorer: a web-based tool for exploratory analysis and visualization of bacterial pan-genomes. Bioinformatics 2022;38:4412–4. 10.1093/bioinformatics/btac504.

[26] Kanehisa M, Sato Y, Morishima K. BlastKOALA and GhostKOALA: KEGG Tools for Functional Characterization of Genome and Metagenome Sequences. J Mol Biol 2016;428:726–31. 10.1016/j.jmb.2015.11.006.

[27] Cantalapiedra CP, Hernández-Plaza A, Letunic I, Bork P, Huerta-Cepas J. eggNOG- mapper v2: Functional Annotation, Orthology Assignments, and Domain Prediction at the Metagenomic Scale. Mol Biol Evol 2021;38:5825–9. 10.1093/molbev/msab293.

[28] Blin K, Shaw S, Augustijn HE, Reitz ZL, Biermann F, Alanjary M, et al. antiSMASH 7.0: new and improved predictions for detection, regulation, chemical structures and visualisation. Nucleic Acids Res 2023;51:W46–50. 10.1093/nar/gkad344.

[29] Ge SX, Jung D, Yao R. ShinyGO: a graphical gene-set enrichment tool for animals and plants. Bioinformatics 2020;36:2628–9. 10.1093/bioinformatics/btz931.

[30] Feng Y, Zou S, Chen H, Yu Y, Ruan Z. BacWGSTdb 2.0: a one-stop repository for bacterial whole-genome sequence typing and source tracking. Nucleic Acids Res 2021;49:D644–50. 10.1093/nar/gkaa821.

[31] Pal C, Bengtsson-Palme J, Rensing C, Kristiansson E, Larsson DGJ. BacMet: antibacterial biocide and metal resistance genes database. Nucleic Acids Res 2014;42:D737–43. 10.1093/nar/gkt1252.

[32] Jia B, Raphenya AR, Alcock B, Waglechner N, Guo P, Tsang KK, et al. CARD 2017: expansion and model-centric curation of the comprehensive antibiotic resistance database. Nucleic Acids Res 2017;45:D566–73. 10.1093/nar/gkw1004.

[33] Zankari E, Hasman H, Cosentino S, Vestergaard M, Rasmussen S, Lund O, et al. Identification of acquired antimicrobial resistance genes. Journal of Antimicrobial Chemotherapy 2012;67:2640–4. 10.1093/jac/dks261.

[34] Feldgarden M, Brover V, Haft DH, Prasad AB, Slotta DJ, Tolstoy I, et al. Validating the AMRFinder Tool and Resistance Gene Database by Using Antimicrobial Resistance Genotype-Phenotype Correlations in a Collection of Isolates. Antimicrob Agents Chemother 2019;63. 10.1128/AAC.00483-19.

[35] Bortolaia V, Kaas RS, Ruppe E, Roberts MC, Schwarz S, Cattoir V, et al. ResFinder 4.0 for predictions of phenotypes from genotypes. Journal of Antimicrobial Chemotherapy 2020;75:3491–500. 10.1093/jac/dkaa345.

[36] Carattoli A, Zankari E, García-Fernández A, Voldby Larsen M, Lund O, Villa L, et al. *In Silico* Detection and Typing of Plasmids using PlasmidFinder and Plasmid Multilocus Sequence Typing. Antimicrob Agents Chemother 2014;58:3895–903. 10.1128/AAC.02412-14.

[37] Brown CL, Mullet J, Hindi F, Stoll JE, Gupta S, Choi M, et al. mobileOG-db: a Manually Curated Database of Protein Families Mediating the Life Cycle of Bacterial Mobile Genetic Elements. Appl Environ Microbiol 2022;88. 10.1128/aem.00991-22.

[38] Wang M, Goh Y-X, Tai C, Wang H, Deng Z, Ou H-Y. VRprofile2: detection of antibiotic resistance-associated mobilome in bacterial pathogens. Nucleic Acids Res 2022;50:W768–73. 10.1093/nar/gkac321.

[39] Vernikos GS, Parkhill J. Interpolated variable order motifs for identification of horizontally acquired DNA: revisiting the *Salmonella* pathogenicity islands. Bioinformatics 2006;22:2196–203. 10.1093/bioinformatics/btl369.

[40] Starikova E V, Tikhonova PO, Prianichnikov NA, Rands CM, Zdobnov EM, Ilina EN, et al. Phigaro: high-throughput prophage sequence annotation. Bioinformatics 2020;36:3882–4. 10.1093/bioinformatics/btaa250.

[41] Wishart DS, Han S, Saha S, Oler E, Peters H, Grant JR, et al. PHASTEST: faster than PHASTER, better than PHAST. Nucleic Acids Res 2023;51:W443–50. 10.1093/nar/gkad382.

[42] Couvin D, Bernheim A, Toffano-Nioche C, Touchon M, Michalik J, Néron B, et al. CRISPRCasFinder, an update of CRISRFinder, includes a portable version, enhanced performance and integrates search for Cas proteins. Nucleic Acids Res 2018;46:W246–51. 10.1093/nar/gky425.

[43] Cosentino S, Voldby Larsen M, Møller Aarestrup F, Lund O. PathogenFinder - Distinguishing Friend from Foe Using Bacterial Whole Genome Sequence Data. PLoS One 2013;8:e77302. 10.1371/journal.pone.0077302.

[44] Kaas RS, Leekitcharoenphon P, Aarestrup FM, Lund O. Solving the Problem of Comparing Whole Bacterial Genomes across Different Sequencing Platforms. PLoS One 2014;9:e104984. 10.1371/journal.pone.0104984.

[45] Letunic I, Bork P. Interactive Tree of Life (iTOL) v6: recent updates to the phylogenetic tree display and annotation tool. Nucleic Acids Res 2024. 10.1093/nar/gkae268.

[46] Liu D, Zhang Y, Fan G, Sun D, Zhang X, Yu Z, et al. IPGA: A handy integrated prokaryotes genome and pan-genome analysis web service. IMeta 2022;1. 10.1002/imt2.55.

[47] HASAN Z. Genome Assembly, Quality Assessment and Annotation of a Multidrug- Resistant, Hypervirulent Klebsiella pneumoniae Isolated from an Infertile Woman Using Spades, Quast, PROKKA and RAST 2025. 10.5281/zenodo.16372706.

[48] Hao L, Yang X, Chen H, Mo Z, Li Y, Wei S, et al. Molecular Characteristics and Quantitative Proteomic Analysis of Klebsiella pneumoniae Strains with Carbapenem and Colistin Resistance. Antibiotics 2022;11:1341. 10.3390/antibiotics11101341.

[49] Fang X, Chen Y, Hu L, Gu S, Zhu J, Hang Y, et al. Modulation of virulence and metabolic profiles in Klebsiella pneumoniae under indole-mediated stress response. Front Cell Infect Microbiol 2025;15. 10.3389/fcimb.2025.1546991.

[50] Zhu J, Wang T, Chen L, Du H. Virulence Factors in Hypervirulent Klebsiella pneumoniae. Front Microbiol 2021;12. 10.3389/fmicb.2021.642484.

[51] Huang J, Zhuang J, Wan L, Liu Y, Du Y, Zhou L, et al. Genomic Analysis and Virulence Assessment of Hypervirulent Klebsiella pneumoniae K16-ST660 in Severe Cervical Necrotizing Fasciitis. International Journal of Medical Microbiology 2024;317:151635. 10.1016/j.ijmm.2024.151635.

[52] Wantuch PL, Knoot CJ, Robinson LS, Vinogradov E, Scott NE, Harding CM, et al. Capsular polysaccharide inhibits vaccine-induced O-antigen antibody binding and function across both classical and hypervirulent K2:O1 strains of Klebsiella pneumoniae. PLoS Pathog 2023;19:e1011367. 10.1371/journal.ppat.1011367.

[53] Hsieh P-F, Lin T-L, Yang F-L, Wu M-C, Pan Y-J, Wu S-H, et al. Lipopolysaccharide O1 Antigen Contributes to the Virulence in Klebsiella pneumoniae Causing Pyogenic Liver Abscess. PLoS One 2012;7:e33155. 10.1371/journal.pone.0033155.

[54] Vats P, Kaur UJ, Rishi P. Heavy metal-induced selection and proliferation of antibiotic resistance: A review. J Appl Microbiol 2022;132:4058–76. 10.1111/jam.15492.

[55] Vinogradov AA, Suga H. Introduction to Thiopeptides: Biological Activity, Biosynthesis, and Strategies for Functional Reprogramming. Cell Chem Biol 2020;27:1032–51. 10.1016/j.chembiol.2020.07.003.

[56] Meletis G. Carbapenem resistance: overview of the problem and future perspectives. Ther Adv Infect Dis 2016;3:15–21. 10.1177/2049936115621709.

[57] uz Zaman T, Aldrees M, Al Johani SM, Alrodayyan M, Aldughashem FA, Balkhy HH. Multi-drug carbapenem-resistant Klebsiella pneumoniae infection carrying the OXA-48 gene and showing variations in outer membrane protein 36 causing an outbreak in a tertiary care hospital in Riyadh, Saudi Arabia. International Journal of Infectious Diseases 2014;28:186–92. 10.1016/j.ijid.2014.05.021.

[58] Ruiz E, Ocampo-Sosa AA, Rezusta A, Revillo MJ, Román E, Torres C, et al. Acquisition of carbapenem resistance in multiresistant Klebsiella pneumoniae strains harbouring bla CTX-M-15, qnrS1 and aac(6′)-Ib-cr genes. J Med Microbiol 2012;61:672–7. 10.1099/jmm.0.038083-0.

[59] Ibisanmi TA, Olowosoke CB, Ayeni TO, Faleti AI. A computational exploration of whole genome sequences of Klebsiella pneumoniae ST16 for beta-lactam resistance and the discovery of NDM-1 resistance gene inhibitor. Inform Med Unlocked 2024;44:101441. 10.1016/j.imu.2023.101441.

[60] Rossolini GM, D’Andrea MM, Mugnaioli C. The spread of CTX-M-type extended- spectrum β-lactamases. Clinical Microbiology and Infection 2008;14:33–41. 10.1111/j.1469-0691.2007.01867.x.

[61] Kawser Z, Sridhar S, Kar S, Habib T, Mukta SA, Azad K, et al. Clinical and genomic characterization of Klebsiella pneumoniae infections in Dhaka, Bangladesh. J Glob Antimicrob Resist 2025;41:52–8. 10.1016/j.jgar.2024.12.016.

[62] Wareth G, Linde J, Hammer P, Pletz MW, Neubauer H, Sprague LD. WGS-Based Phenotyping and Molecular Characterization of the Resistome, Virulome and Plasmid Replicons in Klebsiella pneumoniae Isolates from Powdered Milk Produced in Germany. Microorganisms 2022;10:564. 10.3390/microorganisms10030564.

[63] Sun N, Yang Y, Wang G, Guo L, Liu L, San Z, et al. Whole-genome sequencing of multidrug-resistant Klebsiella pneumoniae with capsular serotype K2 isolates from mink in China. BMC Vet Res 2024;20:356. 10.1186/s12917-024-04222-5.

[64] Kadkhoda H, Gholizadeh P, Ghotaslou R, Pirzadeh T, Ahangarzadeh Rezaee M, Nabizadeh E, et al. Prevalence of the CRISPR-cas system and its association with antibiotic resistance in clinical Klebsiella pneumoniae isolates. BMC Infect Dis 2024;24:554. 10.1186/s12879-024-09451-5.

[65] Li J, Liu Y, Jiang J, Chen F, Zhang N, Kang X, et al. Type I-E* CRISPR-Cas of *Klebsiella pneumoniae* upregulates bacterial virulence by targeting endogenous histidine utilization system. MSphere 2025;10. 10.1128/msphere.00215-25.

[66] Kadkhoda H, Gholizadeh P, Ghotaslou R, Nabizadeh E, Pirzadeh T, Ahangarzadeh Rezaee M, et al. Role of CRISPR-cas system on virulence traits and carbapenem resistance in clinical Klebsiella pneumoniae isolates. Microb Pathog 2025;199:107151. 10.1016/j.micpath.2024.107151.

[67] Shen J, Zhou J, Xu Y, Xiu Z. Prophages contribute to genome plasticity of Klebsiella pneumoniae and may involve the chromosomal integration of ARGs in CG258. Genomics 2020;112:998–1010. 10.1016/j.ygeno.2019.06.016.

[68] Wang F, Wang D, Hou W, Jin Q, Feng J, Zhou D. Evolutionary Diversity of Prophage DNA in Klebsiella pneumoniae Chromosomes. Front Microbiol 2019;10. 10.3389/fmicb.2019.02840.

[69] Struve C, Bojer M, Krogfelt KA. Characterization of Klebsiella pneumoniae type 1 fimbriae by detection of phase variation during colonization and infection and impact on virulence. Infect Immun 2008;76:4055–65. 10.1128/IAI.00494-08.

[70] Khanmohammad KR, Khalili MB, Sadeh M, Talebi AR, Astani A, Shams A, et al. The effect of lipopolysaccharide from uropathogenic Escherichia coli on the immune system, testis tissue, and spermatozoa of BALB/c mice. Clin Exp Reprod Med 2021;48:105–10. 10.5653/cerm.2020.03888.

[71] Muraoka A, Yokoi A, Kajiyama H. Emerging bacterial factors for understanding pathogenesis of endometriosis. IScience 2024;27:108739. 10.1016/j.isci.2023.108739.

